# *Fmr1* mutation reshapes gut microbiome structure, diversity, intestinal barrier integrity, and function in a sex and genotype-dependent manner

**DOI:** 10.64898/2026.03.17.712468

**Authors:** Sabiha Alam, Ciaran A. Shaughnessy, Edralin A. Lucas, Femi Olawale, Babu Fathepure, Elizabeth McCullagh

**Affiliations:** Department of Biology, Oklahoma State University, Stillwater, OK, USA; Department of Nutritional Sciences, Oklahoma State University, Stillwater, OK, USA; Department of Microbiology and Molecular Genetics, Oklahoma State University, Stillwater, OK, USA

**Author notes:** indicates corresponding author.

## Abstract

Fragile X Syndrome (FXS) is the leading monogenic cause of autism spectrum disorder (ASD), caused by mutation in the *Fmr1* gene. In addition to cognitive and behavioral challenges, FXS patients often experience altered gut microbiome-induced gastrointestinal (GI) problems. Evidence suggests that an altered gut microbiome can disrupt mucosal barrier and promote inflammation, leading to impaired intestinal barrier integrity and function. However, the mechanisms by which the gut microbiome alters the barrier integrity and contributes to GI pathology in FXS remain poorly understood. To address this gap, we tested the hypothesis that *Fmr1* mutation reshapes the gut microbiome composition, altering transcriptional markers of intestinal barrier regulation and gut barrier physiology in mice. To test our hypothesis, we used an *Fmr1* knockout (KO) mouse model with wild-type (WT) littermates as controls. First, we performed 16S ribosomal RNA sequencing to characterize gut microbial community structure and diversity across genotypes and sexes. Among the alpha-diversity metrics, only Chao1 showed significant differences across female genotypes in both fecal and cecal contents. Additionally, qRT-PCR analysis of ileal samples revealed reduced barrier and mucosal defense gene expression in female KO and Het mice compared with WT females. Next, we utilized a Ussing chamber assay to test gut epithelial permeability and function. Our physiological data showed that female Het mice had increased transepithelial resistance compared to WT females, indicating a tighter epithelial barrier. Overall, FXS is associated with modest genotype and sex-specific microbiome variation, impaired gut barrier integrity, and altered epithelial barrier function in female mice.

## Introduction

Fragile X Syndrome (FXS) is a neurodevelopmental disorder caused by a CGG trinucleotide repeat expansion in the 5′ untranslated region of the *FMR1* (Fragile X Messenger Ribonucleoprotein 1) gene on the X chromosome (1–4). Mutation in the *FMR1* gene leads to reduction or elimination of the protein product FMRP (Fragile X Messenger Ribonucleoprotein (5–7). FMRP is a conserved RNA-binding protein that plays a critical role in the translation of many aspects of target mRNAs involved in neuronal development and cellular function (8, 9).

In addition to well-known cognitive and behavioral challenges, people with FXS also experience serious gastrointestinal (GI) complications. However, the mechanisms of gut-specific phenotypes are not well understood. Previous reports demonstrate that individuals with FXS frequently experience GI disturbances, including constipation, diarrhea, irritable bowel syndrome, and gastroesophageal reflux (10, 11). Recent work has identified alterations in intestinal microbial composition and predicted metabolic pathways in male *Fmr1* KO2 mice, highlighting the potential relevance of microbiome-associated GI disorders in FXS (12). A growing body of literature suggests a central role for the gut microbiome structure in maintaining epithelial barrier function. Gut dysbiosis leads to increased intestinal permeability and mucosal inflammation across multiple disease contexts, including ASDs (13–18). This body of work therefore warrants further investigation into whether GI dysfunction in FXS might also stem from microbiome-associated disruptions in intestinal barrier homeostasis.

Sex differences represent a crucial yet underexplored area in the FXS and ASD research fields, although both conditions are diagnosed more frequently in males (19). Females often exhibit distinct and heterogeneous phenotypes due to X-linked genetic regulation, mosaic *Fmr1* expression, and sex-dependent hormonal and immune influences (20–22). In parallel, gut microbiome structure, host–microbe interactions, and intestinal barrier physiology are increasingly recognized as sex-modulated processes (23–25). GI manifestations in ASD and FXS may therefore differ between males and females. Importantly, in our experiments, inclusion of heterozygous (*Fmr1*^*+/-*^) females introduces an intermediate genetic context that models mosaic *Fmr1* expression observed in many affected human females and enables direct assessment of *Fmr1* gene-dosage– dependent effects. This experimental approach permits us to evaluate how mosaicism in FXS influences gut microbial community, epithelial barrier integrity, and physiology, supporting a clearer interpretation of genotype- and sex-specific gut barrier homeostasis.

In this study, we integrated host-microbiota-epithelial barrier interactions in both male and female FXS mice. Together, this is first ever systematic sex inclusive evaluation of gut-barrier-associated features using an FXS mouse model and establishes a foundation for identifying microbial and epithelial signatures that may contribute to establishing GI phenotypes and potential biomarkers in FXS.

## Methodology

### Animals

All animal procedures were conducted in accordance with the National Institutes of Health Guide for the Care and Use of Laboratory Animals and complied with ARRIVE guidelines. Experimental protocols were reviewed and approved by the Institutional Animal Care and Use Committee (IACUC) at Oklahoma State University. All experiments were conducted using *Fmr1* KO mice (JAX strain B6.129P2-Fmr1<sup>tm1Cgr</sup>/J, stock #003025) maintained on a wild-type background, (JAX, stock #000664) which we used as controls. Experimental groups included hemizygous *Fmr1* KO males, homozygous KO females, and heterozygous females. All mice were originally sourced from The Jackson Laboratory and subsequently bred at Oklahoma State University (26). Animals used in these experiments were bred from mixed and single genotype matings to produce heterozygotes and littermate controls, as well as to maintain breeding lines. Figure legends list the number of animals used in each experiment. All animals used in this study were between 60 and 120 days old. All efforts were made to minimize animal suffering and to reduce the number of animals used.

### 16S rRNA sequencing for microbiome structure and microbial diversity analysis Sample collection

After sacrificing mice with isoflurane, we collected fecal (n=30) and cecal (n=30) contents, freeze-dried them at 80 °C, until they were processed for DNA extraction. Cecal contents were collected by opening the cecum under sterile conditions, transferring the luminal contents into sterile tubes, and immediately freezing the samples at −80 °C. Microbiome analyses were performed across the same five experimental groups, with n = 6 per group for both cecal and fecal materials.

### Library construction, sequencing, and bioinformatics analysis

We extracted total genomic DNA from each sample using the QIAamp Power-Fecal DNA Kit (Qiagen, Germantown, MD, USA) following the manufacturer’s protocol. 16S rRNA gene regions were amplified using primers targeting conserved sequences with Illumina sequencing adapters attached. We amplified the V3–V4 hypervariable regions of the bacterial 16S rRNA gene with universal primers 388 F (5’-ACTCCTACGGGAGGCAGCA-3’) and 806 R (5’-GGACTACHVGGGTWTCTAAT-3’). Polymerase chain reaction (PCR) was performed under the following thermal cycling conditions: initial denaturation at 95 °C for 5 minutes; followed by 20 cycles of denaturation at 95 °C for 30 seconds, annealing at 50 °C for 30 seconds, and extension at 72 °C for 40 seconds; with a final extension at 72 °C for 7 minutes. The resulting amplicon products were purified using the Omega DNA purification kit (Omega Inc., Norcross, GA, USA), quantified and homogenized to get a sequencing library. Library QC was performed for constructing libraries, and qualified libraries were sequenced on the Illumina Novaseq 6000 platform to generate paired-end reads at Biomarker Technologies (BMKGENE USA Inc., Durham, NC). Raw image data were converted to FASTQ format by base calling. Quality control of raw reads was performed using Trimmomatic v0.33 (7), for quality trimming and Cutadapt v1.9.1 for primer removal (28). High-quality reads were processed in QIIME2 (29), using the DADA2 algorithm for denoising (30), chimera removal, and generation of amplicon sequence variants (ASVs), which provides single-nucleotide resolution (31).

Taxonomic classification of ASVs was performed using a Bayesian classifier and BLAST against the SILVA 138 reference database (32). Features with read counts <2 were filtered out. Taxonomic compositions were summarized at multiple ranks using QIIME2 feature-table and taxonomy artifacts. Relative abundances were calculated and collapsed at the genus level. The top 20 genera were visualized as stacked bar plots, along with alpha-diversity (Shannon, Simpson and Chao1) and beta-diversity (Bray–Curtis dissimilarity with NMDS-coordination, which were assessed in QIIME2 (33) and visualized in the BMKCloud platform (34). ALDEx2 differential abundance analysis and comparisons of bacterial genera across male and female genotypes were performed in RStudio using the ALDEx2 (35), ggplot2, dplyr, tidyr, and tibble packages (36, 37).

### Gut barrier integrity profiling

#### Sample collection

Experimental mice were fasted for at least 3 hours and then euthanized with a mixture of ketamine and xylazine (100 mg/kg and 10 mg/kg). After collecting their ilea, we flushed those tissues with ice-cold sterile saline, then preserved them in liquid nitrogen followed by -80 °C until mRNA extraction.

We extracted ileal total RNA using a Trizol RNA isolation reagent (Thermo Fisher Scientific, Waltham, MA, USA). The relative abundance of genes (*Tjp1, Ocln, Cldn2, Plvap, Muc2*, Reg*3g, Alpi and Tnf*) was quantified by quantitative reverse transcription-polymerase chain reaction (qRT-PCR). qRT-PCR was performed using SYBR Green chemistry on an ABI 7900HT sequence detection system equipped with SDS 2.4 software (Applied Biosystems). Relative transcript levels were quantified using the 2^−ΔΔCt method, employing GAPDH as the endogenous reference gene.

#### Gut barrier physiology

To evaluate intestinal epithelial physiology and barrier function in FXS mice, we performed *ex vivo* Ussing chamber electrophysiological experiments on intestinal tissues. Freshly dissected distal ileum segments were gently flushed with PBS to remove luminal contents, cut longitudinally, and opened to create a full-thickness two-dimensional tissue plane to be mounted in the Ussing chamber baths (P2300, Physiologic Instruments, Inc., Venice, FL, USA). The tissue was mounted flat on an EasyMount Ussing Chamber Slider with an exposed tissue area of 0.04 cm^2^ and placed between mucosal and serosal baths. The baths contained identical Ringer’s solution: 120 mM NaCl, 10 mM D-glucose, 3.3 mM KH_2_PO_4_, 0.83 mM K_2_HPO_4_, 1.2 mM MgCl_2_, 1.2 mM CaCl_2_, pH 7.4. Mounted tissues were short-circuited via a voltage-clamp system (VCC MC8-8S, Physio Instruments, Inc.) and maintained for up to 2 hr at 37 °C, saturated with of 95% O_2_/5% CO_2_ gas mixture. Once epithelial preparations stabilized under these conditions, values for short-circuit current (I_sc_), voltage (V_t_), conductance (G_t_), and electrical resistance (TER) were obtained for analysis. These baseline measurements were used to provide indirect electrical assessments of epithelial barrier properties of the native gut tissue. It is important to note that measurements do not distinguish pore versus subepithelial contributions.

#### Statistical analyses

All statistical analyses on data derived from gut barrier integrity and function experiments were performed in R (version 4.3.2) using RStudio (version 2023.06.1). Normality of each dataset was evaluated with the Shapiro–Wilk test (stats package), and homogeneity of variances was assessed using Levene’s test (car package). Depending on distributional properties, for parametric data, to assess group differences, one-way ANOVA with Tukey post hoc tests (stats packages) was applied. Parallelly, for nonparametric data, Kruskal–Wallis tests followed by Dunn’s post hoc comparisons (FSA package) were performed. Between-group comparisons, Student’s t-test, or Welch’s t-test were applied as appropriate. When assumptions were violated, Wilcoxon rank-sum tests (coin package) were used (37). Statistical significance was defined as *p* < 0.05.

## Results

### Characterization of the gut microbiome structure and diversity

#### Genotype-associated variation in microbial communities of the cecal contents in male mice

Microbial community composition and diversity can modulate FXS-associated symptoms in both clinical and preclinical settings (38–40). In our study, we considered the top 20 genera which revealed broadly similar community structures between male WT and KO groups (Fig. 1A), indicating genotype-associated effects were distributed across multiple taxa in cecal contents. Alpha-diversity metrics indicated no significant differences in overall microbial richness or evenness between WT and KO males. The Simpson’s index showed comparable diversity across genotypes (Fig. 1B, p = 0.39), and neither Chao1 (p = 0.59, Fig. 1C) nor Shannon diversity (p = 0.065, Fig. 1D) differed significantly, indicating preservation within-sample-based microbiome diversity despite *Fmr1* deficiency in male mice. In contrast, β-diversity analysis revealed partial separation of microbial communities by genotype. NMDS ordination based on Bray–Curtis dissimilarity showed overlapping yet distinguishable clustering of WT and KO samples (Fig. 1E). This pattern was supported by PERMANOVA (R^2^ = 0.235, p = 0.003) analysis, indicating a significant but modest genotype effect on community composition. Together, these findings suggest that *Fmr1* mutation in male mice is associated with subtle but detectable shifts in gut microbiome structure without major alterations in overall alpha diversity. In this study, we also considered fecal contents; we found no significant genotype-associated differences between WT and KO males, with comparable alpha diversity (Simpson p = 0.94, Chao1 p = 0.82, Shannon p = 0.94, supplementary Figs. 1C-1D) and no separation in β-diversity (R^2^ = 0.078, p = 0.581), despite minor compositional variation (supplementary Fig. 1E).

**Figure 1.**
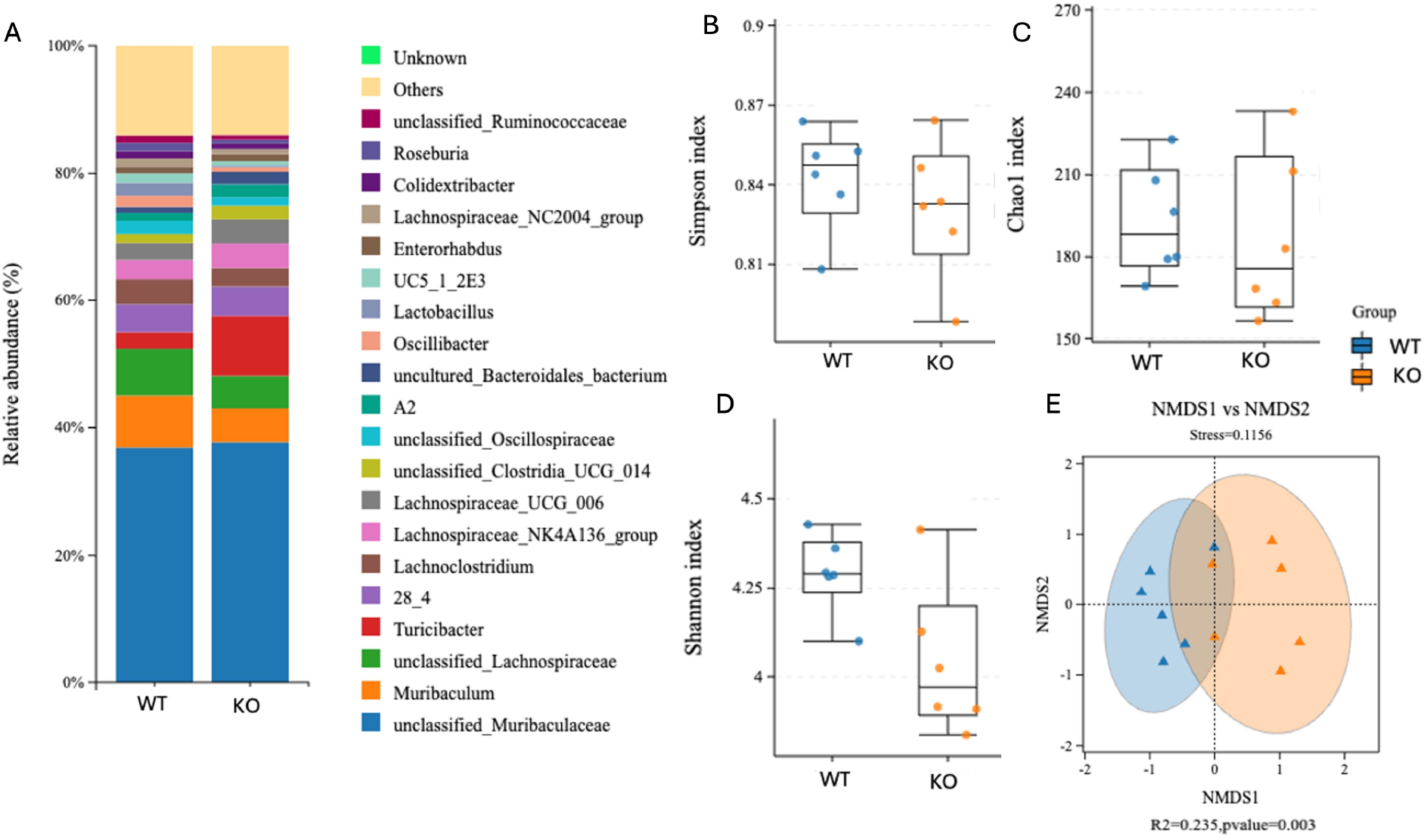
Male gut microbiome composition and microbial diversity in WT and KO cecal contents. (A) Relative abundance of the top 20 bacterial genera in cecal contents from WT and KO male mice. (A) Genus-level microbial composition displayed as stacked bar plots showing relative abundance; low-abundance taxa are grouped as “Others,” and unassigned taxa are denoted as “Unknown.” (B–D) Alpha-diversity metrics comparing WT and KO males, including the Simpson index (B), which reflects microbial species evenness and dominance, the Chao1 index (C), an estimator of species richness, and the Shannon diversity index (D), which integrates both species richness and evenness. Statistical significance was assessed using the Mann–Whitney U test. Each point represents an individual animal, and boxplots depict the median and interquartile range. (E) β-diversity analysis based on Bray–Curtis dissimilarity at the species level, visualized by non-metric multidimensional scaling (NMDS). Group differences were evaluated using PERMANOVA, with shaded ellipses representing 95% confidence intervals for each genotype. Data are derived from six independent biological replicates per group (n = 6 per genotype).

We also performed differential abundance analysis using ALDEx2, applying Dirichlet Monte Carlo sampling and CLR transformation to account for compositional microbiome data. Statistical significance was assessed by Welch’s t-test for pairwise comparisons in male study groups and the Kruskal–Wallis test for multi-group comparisons in female study groups, with p-values corrected for multiple testing using the Benjamini–Hochberg false discovery rate (FDR). However, we did not find any genera that remained significant after FDR correction across experimental group comparisons (supplementary Figs. 9A-9D).

#### Genotype-associated variation of microbial communities in the cecal contents of female mice

Gut microbial community composition showed modest genotype-associated differences across WT, Het, and KO female mice. We analyzed the top nine bacterial genera based on their relative abundance. However, none of those top nine genera showed a significant difference across female genotypes. In this study, alpha diversity metrics revealed selective genotype effects in female mice. For example, Chao1 index differed modestly across genotypes, with Het mice generally exhibiting higher diversity or species richness relative to WT (p = 0.0022), while KO mice did not show greater variability compared to WT females (p = 0.48) and Hets (p = 0.13) in Fig. 2C. For Shannon diversity index, WT females compared to Het females showed no significant difference (p = 0.39, Fig. 2D) and the same pattern was exhibited when comparing Hets with KO females (p = 0.94, Fig. 2D). Simpson index, which is also an alpha diversity parameter, and usually indicates dominance structure across species, also showed no significant differences across female genotypes (Fig. 2B, WT vs Het, p = 0.82, WT vs KO, p = 0.82 and for WT vs KO, p = 1). However, female fecal samples showed higher Chao1 index in Hets compared to WT females (p = 0.0087, supplementary Fig. 2C), while other parameters did not show any significance alpha diversity in fecal female study groups (supplementary Figs. 2B and 2D). In contrast, β-diversity analysis in female fecal contents demonstrated a significant but partial separation of microbial communities by genotype (R^2^ = 0.21, p = 0.019, supplementary Fig. 2E).

**Figure 2.**
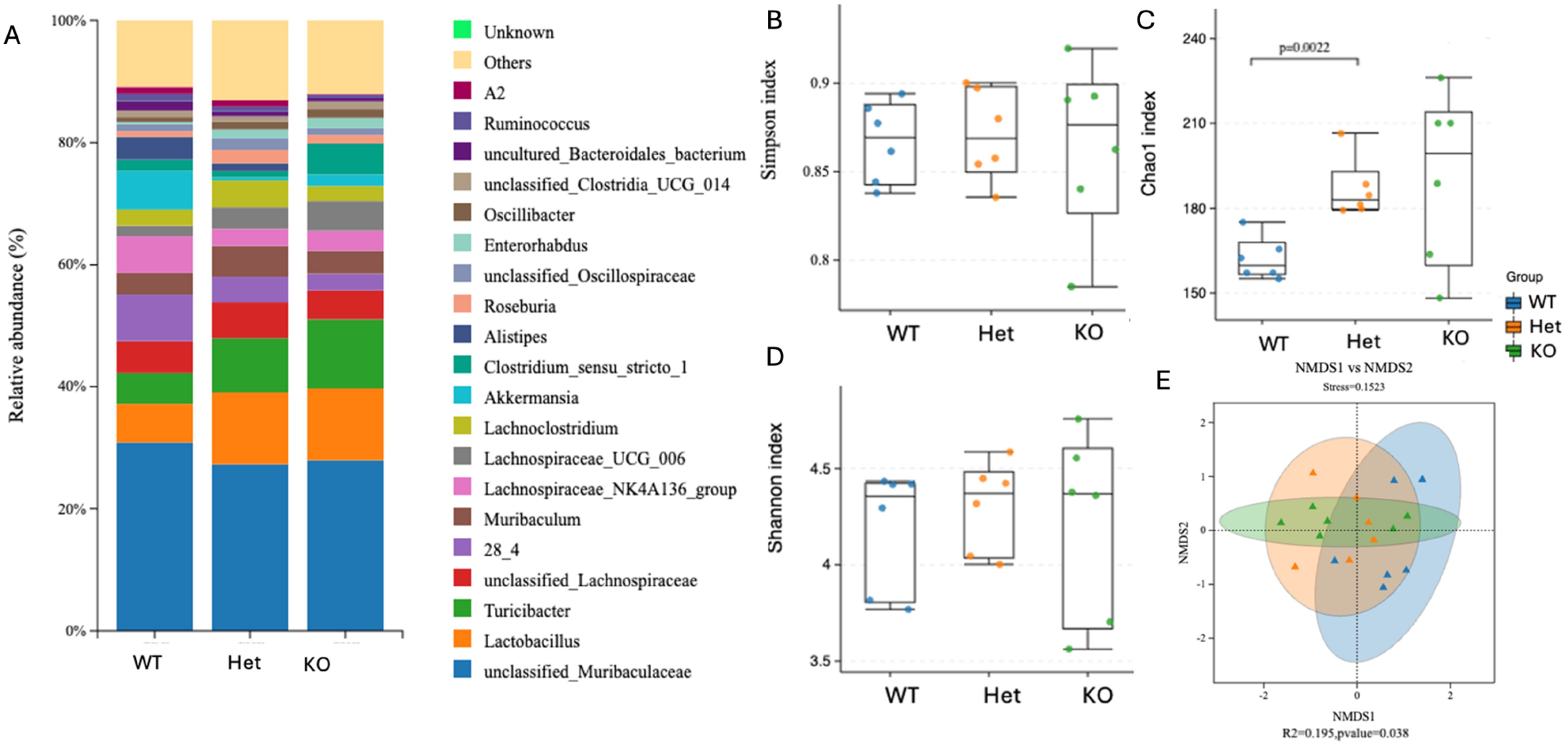
Female cecal gut microbiome composition and microbial diversity among WT, Het and KO mice. (A) Relative abundance of the top 20 bacterial genera in cecal contents across female WT, Het, and KO mice. (A) Genus-level microbial composition displayed as stacked bar plots showing relative abundance; low-abundance taxa are grouped as “Others,” and unassigned taxa are denoted as “Unknown.” (B–D) Alpha-diversity metrics comparing WT, Het, and KO females, including the Simpson index (B), which reflects microbial species evenness and dominance, the Chao1 index (C), an estimator of species richness, and the Shannon diversity index (D), which integrates both richness and evenness. Statistical significance was assessed using the Mann–Whitney U test. Each data point represents an individual animal, and box plots depict the median and interquartile range. (E) β-diversity analysis based on Bray–Curtis dissimilarity at the species level, visualized by non-metric multidimensional scaling (NMDS). Group differences were evaluated using PERMANOVA, with shaded ellipses representing 95% confidence intervals for each genotype. Data are derived from six independent biological replicates per group (n = 6 per genotype).

### Effects of sex difference within genotype on gut microbiome structure and microbial diversity in cecal and fecal samples

#### Cecal Contents

We first evaluated sex-specific differences in gut microbial communities in cecal contents of WT mice. Taxonomic profiling indicated sex-associated shifts in microbial composition (supplementary Fig. 3A). Although overall community diversity did not differ between WT males and females based on Simpson (p = 0.18) or Shannon indices (p = 0.82), Chao1 analysis revealed significantly greater species richness in males (p = 0.0043, supplementary Figs. 3B–3D). Consistent with this observation, our study revealed that microbiome community structure had a significant separation between male and female microbial communities (R^2^ = 0.31, p = 0.001, supplementary Fig. 3E).

A similar pattern was observed in cecal KO male and KO female mice. While Simpson (p = 0.18) and Shannon diversity (p = 0.59) did not differ between KO males and KO females, Chao1 richness remained significantly higher in males (p = 0.0043, supplementary Figs. 4B–4D). Correspondingly, β-diversity analysis revealed significant separation between KO male and KO female microbial communities (R^2^ = 21, p = 0.001, supplementary Fig. 4E). We next examined whether the mosaic genotype effect produces distinct microbial patterns relative to fully deficient KO mice by comparing KO males with Het females. In this comparison, Het females exhibited significantly higher Simpson diversity than KO males (p = 0.026), whereas Chao1 richness did not differ between groups (p = 0.7). The Shannon index showed a trend toward higher diversity in Het females but did not reach statistical significance (p = 0.065, supplementary Figs. 5B–5D). Despite these modest differences in alpha diversity, β-diversity analysis demonstrated a clear separation between KO male and Het female microbial communities (R^2^ = 0.248, p = 0.001, supplementary Fig. 5E), indicating distinct microbial community structures between the two groups.

#### Fecal Contents

To determine whether sex influences gut microbial structure in the feces, we first examined male– female differences within WT mice. Taxonomic profiling suggested sex-associated shifts in microbial composition (supplementary Fig. 6A). While overall alpha diversity did not differ between males and females based on Simpson (p = 0.31) or Shannon indices (p = 0.18), Chao1 richness was significantly higher in males (p = 0.0087, supplementary Figs. 6B–6D). Consistent with these observations, β-diversity analysis revealed a significant separation between male and female microbial communities (R^2^ = 0.252, p = 0.008, supplementary Fig. 6E), indicating sex-dependent differences in community structure.

We next assessed whether similar sex-associated patterns are maintained between KO males and KO females. In KO male mice, alpha diversity indices again did not differ significantly between males and females (Simpson p = 0.48, Chao1 p = 0.093, Shannon p = 0.24, supplementary Figs. 7B–7D). However, β-diversity analysis still showed a significant separation between KO male and female microbial communities (R^2^ = 0.175, p = 0.035, supplementary Fig. 7E), suggesting that sex-dependent differences in overall microbial community structure persist even in the absence of FMRP. In fecal KO male vs Het female comparison, we found that alpha diversity did not differ between groups based on Simpson (p = 0.48), Chao1 richness (p = 0.59), or Shannon indices (p = 0.31, supplementary Figs. 8B–8D). Consistent with these findings, β-diversity analysis did not reveal significant separation between KO male and Het female microbial communities in feces (R^2^ = 0.071, p = 0.586, supplementary Fig. 8E). Overall, these results indicate that sex is a major determinant of gut microbial community structure in both WT and *Fmr1* KO mice, whereas the mosaic genotype in Het females produces more modest effects on microbial diversity and composition.

### Deletion of Fmr1 alters gut barrier integrity markers in mice

Using RT-qPCR, we examined gene expression of intestinal samples to assess epithelial barrier integrity markers including *Tjp1, Ocln, Cldn2, Plvap, Muc2, Reg3g, Alpi*, and *Tnf*.

### Genotype-dependent gut barrier integrity profiling in male mice

Expression levels of *Tjp1, Ocln, Cldn2, Plvap, Muc2, Reg3g, Alpi*, as well as *Tnf* showed no significant differences between genotypes (all p > 0.05) in male mice (Fig. 3). Overall, *Fmr1* loss does not alter gut barrier integrity or inflammatory status in male mice (Figs. 3A–3H).

**Figure 3.**
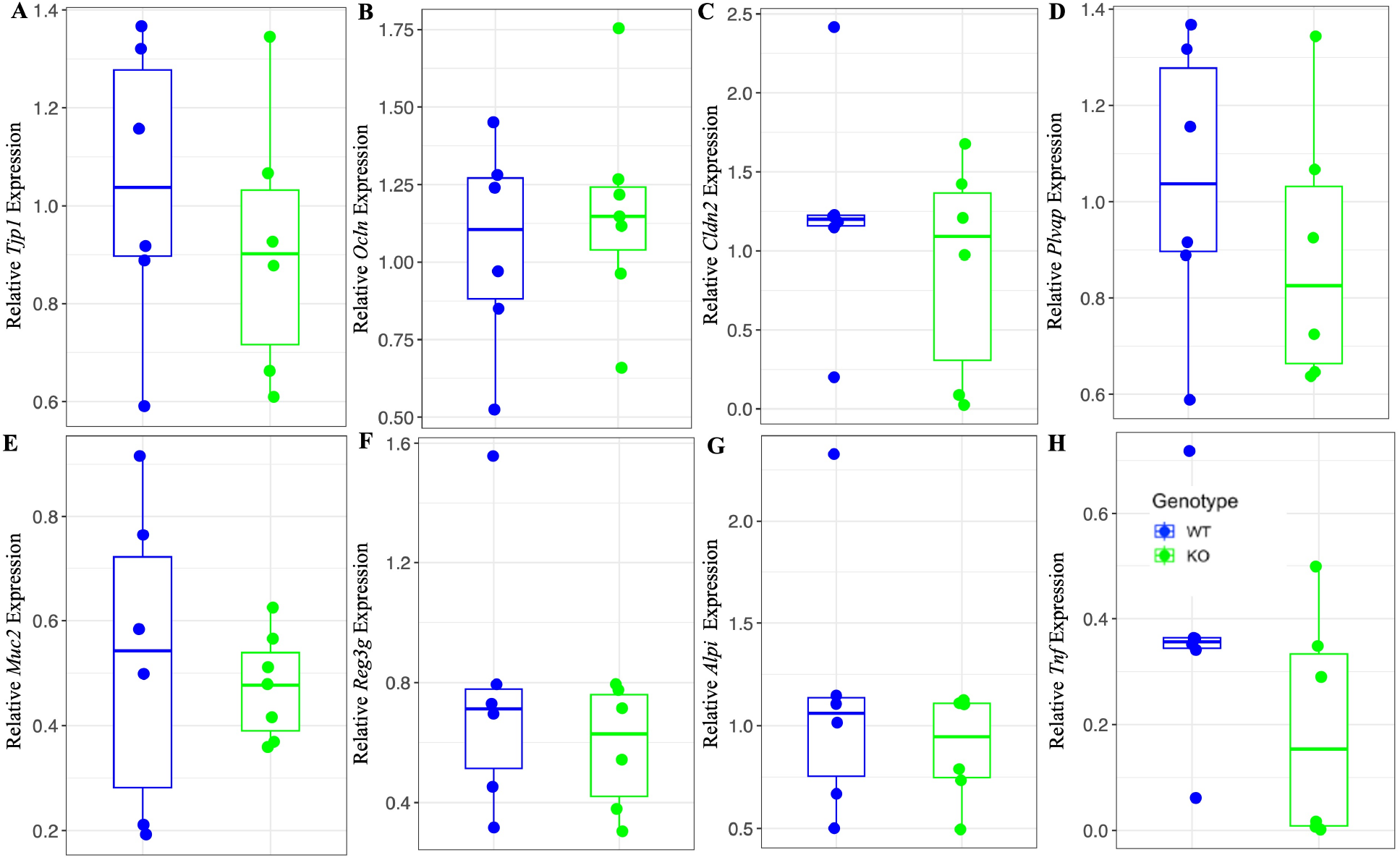
Transcriptional markers of intestinal barrier structure, mucosal defense, and inflammatory tone in male WT and KO mice. Relative expression of genes collectively involved in maintaining the intestinal barrier in male mice (A) *Tjp1* and (B) *Ocln*, core tight junction–associated proteins (C) *Cldn2*, a pore-forming tight junction component (D) *Plvap*, an endothelial permeability marker (E) *Muc2*, a mucus layer structural component (F) *Reg3g*, an epithelial anti-microbial peptide (G) *Alpi*, an anti-inflammatory epithelial enzyme and (H) *Tnf*, a pro-inflammatory cytokine. Box-and-whisker plots depict the median (center line), interquartile range (box), and full data range (whiskers), with individual points representing independent biological replicates. Statistical analyses were selected based on Shapiro– Wilk normality and homogeneity of variance testing: unpaired two-tailed Student’s t tests were applied for panels A, B, D, and H; Welch’s t test for panel C; and Mann–Whitney U tests for panels E, F, and G. Each group included 5–7 independent biological replicates (n = 5–7 per group). Exact p-values are indicated in the respective panels.

### Genotype-dependent gut barrier integrity profiling in female mice

Comparisons among WT, Het, and KO female mice revealed significantly reduced expression of epithelial tight junction markers (Figs. 4A-4C). Specifically, in Fig. 4A, *Tjp1* expression was decreased in KO females compared with WT (p = 0.0252) and in Het females compared with WT (p = 0.034). *Ocln* expression was also significantly reduced in KO females relative to WT controls (p = 0.025, Fig. 4B). Additionally, the endothelial tight junction marker *Plvap* showed reduced expression in KO females compared with WT (p = 0.03) and in Het females compared with KO (p = 0.033, Fig. 4D). Collectively, these findings indicate a genotype-dependent disruption of intestinal barrier components in female *Fmr1*-deficient mice.

**Figure 4.**
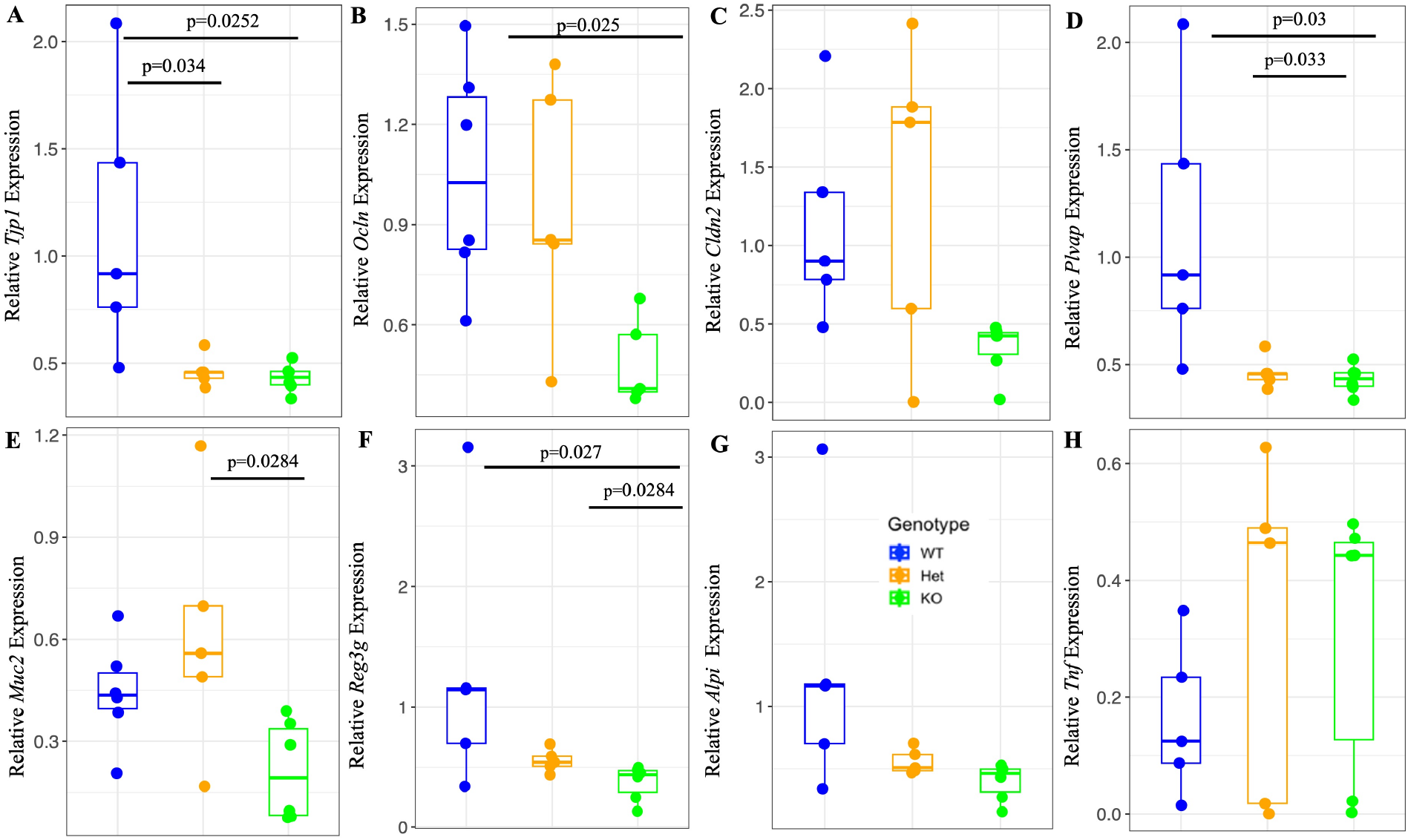
Transcriptional markers of intestinal barrier structure, mucosal defense, and inflammatory tone among female WT, Het and KO mice. Relative expression of genes collectively involved in maintaining the intestinal barrier integrity in wild-type (WT), heterozygous (Het), and knockout (KO) female mice. (A) *Tjp1*; (B) *Ocln;* (C*) Cldn2*; (D) *Plvap*; (E) *Muc2*; (F) *Reg3g*; (G) *Alp*i; and (H) *Tnf*. Box-and-whisker plots depict the median (center line), interquartile range (box), and full data range (whiskers), with individual points representing independent biological replicates. Statistical analyses were selected based on Shapiro–Wilk normality and homogeneity of variance testing. One-way ANOVA was applied for panel B, while Kruskal–Wallis tests were used for panels A and C–H. When overall significance was detected, Tukey’s post hoc test (ANOVA) or Dunn’s post hoc test (Kruskal–Wallis) was applied, as indicated, based on normality tests. Each group included 5–7 independent biological replicates (n = 5–7 per group). Exact p-values are indicated in the respective panels.

In contrast, *Muc2* expression was significantly lower in KO compared with Het (Fig. 4E, p = 0.0284). The *Reg3g* expression was significantly different in between WT vs KO (p = 0.027), and in Hets vs KO mice (p = 0.0284, Fig. 4F). Expression of *Cldn2* did not differ significantly across genotypes, although trend-level changes were observed for *Cldn2* (p = 0.0604 for WT vs KO, p = 0.0687 for Het vs KO, Fig. 4C). Also, no significant differences were observed among WT, Het, and KO groups for *Alpi* (p = 0.068, Fig. 4G) and *Tnf* (p = 0.591, Fig. 4H). Overall, *Fmr1* loss in females is associated with selective impairment of epithelial barrier and anti-microbial gene expression, without a marked pro-inflammatory response across their genotypes.

#### Sex-specific differences within genotype in transcriptional regulators of gut barrier integrity

We measured sex specific influence of transcriptional regulators within genotype across male and female mice. In WT mice, we did not observe any significant sex-associated differences in the expression of gut barrier–related genes (*Tjp1, Ocln, Cldn2, Plvap, Muc2, Reg3g, Alpi, Tnf*) between males and females (all p > 0.05, supplementary Figs. 10A-10H). Although *Tnf* showed a modest trend toward higher expression in males (p = 0.08, supplementary Fig. 10H).

In contrast, KO mice exhibited significant sex-dependent differences in several barrier-associated genes. KO males showed higher expression of *Tjp1* compared with KO females (p = 0.0065, supplementary Fig. 11A) and the same pattern was observed in *Alpi* (p = 0.0023, supplementary Fig. 11G), while other genes did not differ significantly between full FXS KO male and females. Similarly, when KO males were compared with Het females, KO males displayed significantly higher expression of *Tjp1* (p = 0.0042, supplementary Fig. 12A), *Plvap* (p = 0.0043, supplementary Fig. 12D), and *Alpi* (p = 0.0032, supplementary Fig. 12G), compared with Het females, whereas the remaining genes showed no significant differences. Overall, these results indicate that *Fmr1* mutation advances sex and genotype dependent alterations in key epithelial barrier integrity maintaining genes, particularly *Tjp1, Plvap*, and *Alpi*.

### Gut physiology analysis

#### Gut barrier physiology status in male mice across genotypes

Our data showed a significant increase in I_sc_ in KO males compared with WT males (p = 0.037), indicating an altered active ion transport in KO male mice. In contrast, other epithelial barrier markers such as V_t_, G_t_, and TER, did not differ across genotypes in male mice (Fig. 5B-5D). Together with the *Tjp1, Ocln*, and *Cldn2* expression data, these data may indicate that *Fmr1* loss in male mice selectively affects epithelial electrogenic ion transport processes while having minimal effect on barrier integrity.

**Figure 5.**
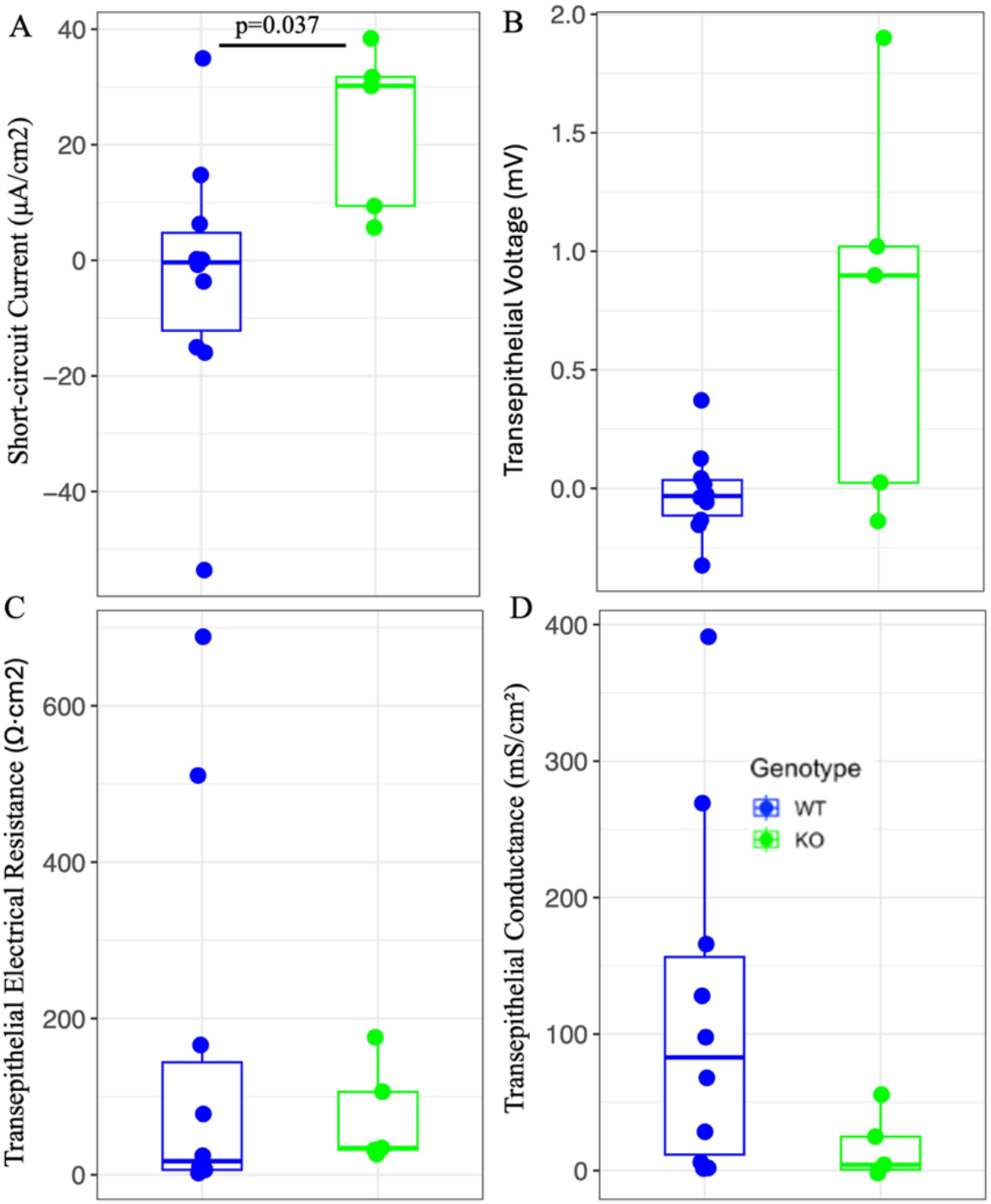
Intestinal paracellular barrier physiology in male WT and KO mice. *Ex vivo* assessment of intestinal epithelial barrier function in male WT and KO mice using Ussing chamber electrophysiology. (A) Short-circuit Current (I_sc_), (B) Transepithelial voltage (V_t_), (C) Transepithelial electrical resistance (TER), and (D) transepithelial conductance (G_t_). Box-and-whisker plots depict the median (center line), interquartile range (box), and full data range (whiskers), with individual points representing independent biological replicates. Statistical analyses were selected based on Shapiro–Wilk normality and homogeneity of variance testing: unpaired two-tailed Student’s t tests were applied for panels A–C, and a Mann–Whitney U test was used for panel D. Each group included 5–9 independent biological replicates (n = 5–9 per group). Exact p-values are indicated in the respective panels.

### Gut barrier physiology status in female mice across genotypes

Ussing chamber analyses showed no genotype-dependent differences in I_sc_ or V_t_ (Figs. 6A and 6B) for female mice. In contrast, genotype-dependent differences were detected for TER and G_t_ for female mice. Het females exhibited significantly increased TER (p = 0.008, Fig. 6C) and reduced G_t_ compared with WT (p = 0.008, Fig. 6D), indicating enhanced barrier tightness. KO females also showed a trend of elevated TER compared to WT females, although this was not significant (p = 0.1). No differences in G_t_ were detected between KO and WT females (p = 0.397). Overall, these data indicate genotype-dependent modulation of gut barrier function exists in females without accompanying changes in electrogenic ion transport.

**Figure 6.**
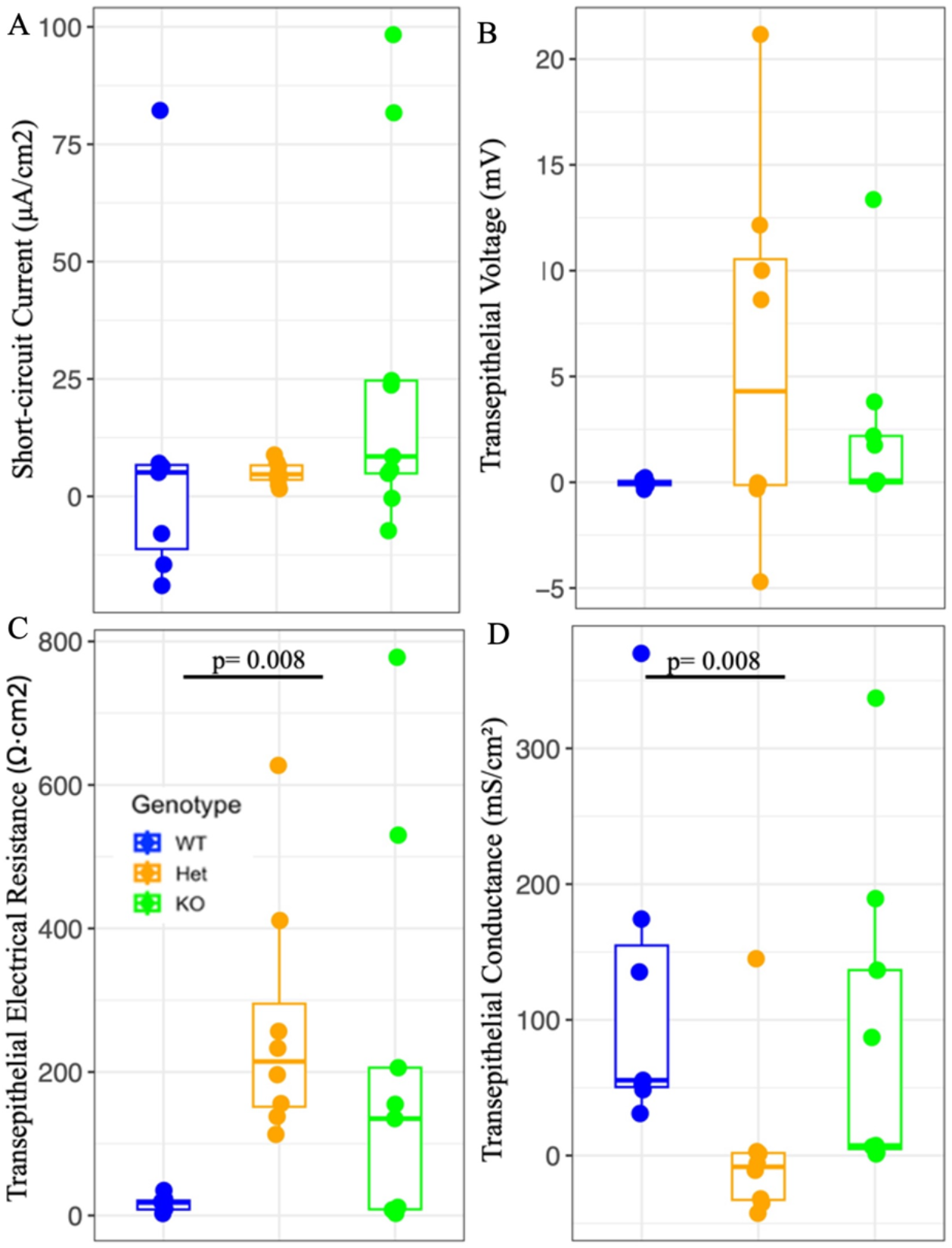
Intestinal paracellular barrier physiology among female WT, Het and KO mice. *Ex vivo* assessment of intestinal epithelial barrier function in female WT, Het, and KO mice using Ussing chamber electrophysiology. (A) Short-circuit current (I_sc_), (B) transepithelial voltage (V_t_), (C) transepithelial electrical resistance (TER), and (D) transepithelial conductance (G_t_). Box-and-whisker plots depict the median (center line), interquartile range (box), and full data range (whiskers), with individual points representing independent biological replicates. Statistical analyses were selected based on Shapiro–Wilk normality and homogeneity of variance testing. Kruskal–Wallis tests were applied for panels A–D, followed by Dunn’s post hoc tests when overall significance was detected. Each group included 5–9 independent biological replicates (n = 5–9 per group). Exact p-values are indicated in the respective panels.

### Sex-specific differences within genotype in gut barrier physiology

Using the Ussing chamber assay, we evaluated sex-associated differences within genotype in intestinal epithelial barrier function. Comparing WT male and female mice, we did not observe significant sex-associated differences in short-circuit current (Isc), transepithelial voltage, transepithelial electrical resistance (TER), or epithelial conductance (G_t_) (all p > 0.05, supplementary Figs. 13A-13D). Similarly, we found no significant differences between KO males and KO females in any electrophysiological parameter (supplementary Figs. 14A-14D). However, when we compared KO males with Het females, significant gut barrier related functional differences emerged. We observed higher Isc in KO males compared with Het females (p = 0.011, supplementary Fig. 15A), whereas TER was significantly higher in Het females (p = 0.049, supplementary Fig. 15C). In contrast, V_t_ and G_t_ did not differ significantly between Het females and KO males. Together, these results suggest that mosaic *Fmr1* expression in Het females is associated with improved epithelial barrier integrity relative to KO males.

## Discussion

This study embodies a systems-level characterization of gut microbial ecology, defining microbiome structure and diversity along with intestinal barrier integrity and function in a FXS mouse model. This holistic characterization demonstrates how *Fmr1* deficiency drives the remodeling of the overall gut ecosystem, extending beyond microbial composition to altered epithelial barrier niches in FXS mice. Notably, female Het mice showed the most pronounced gut-associated phenotypes, including altered microbial diversity and epithelial barrier remodeling. These findings highlight a previously underappreciated role of the *Fmr1* mutation in sex-specific host–gut–microbiome–epithelial barrier interactions. These findings also signify *Fmr1* as a key regulator at the interface of the gut microbiome and intestinal physiology, suggesting that GI dysfunction observed in FXS arises from coordinated disruptions in microbial–epithelial crosstalk rather than from microbial dysbiosis alone.

In this study, analysis of the gut microbiome revealed divergence in microbial composition in both male and female mice across genotypes. This pattern aligns with increasing evidence that neurodevelopmental disorders including ASD-associated syndromes, are accompanied by alterations in gut microbial community structure (3, 41). Altered microbial diversity observed may reflect disruptions in host–microbe signaling pathways particularly those regulating mucosal immune tone, metabolic functions, or epithelial differentiation (40, 42–44).These findings further support the concept that gut dysbiosis is not merely a secondary phenotype of FXS but may contribute directly to intestinal phenotypes. To further evaluate genotype-associated microbial variation, we examined genus-level relative abundance differences for the nine most abundant genera across experimental groups such as-*Lachnoclostridium, Clostridium sensu stricto 1, Lachnospir*aceae_NK4A136_group, Lachnospiraceae_UCG_006, *Lactobacillus, Muribaculum, Turicibacter*, unclassified_Lachnospiraceae, and unclassified_Muribaculaceae. Among all our 5 experimental groups, cecal samples in males across genotypes showed significant differences in 2 of those 9 taxa such as-*Lachnoclosdrium* and *Lactobacillus*. A systematic review reported that *Lactobacillus* supplementation with fecal microbiota transplantation from healthy donors improved constipation and behavioral symptoms in children with ASD. Moreover, perinatal administration of *Lactobacillus* prevented Asperger- and ADHD-like symptoms in adolescence (45). But the role of *Lachnoclostridium* in ASD remains inconsistent across studies. While some reports have found reduced abundance of this genus in ASD (46), others have observed increased levels in ASD populations (47). *Muribaculum* and *Turicibacter* showed nominal differences (p < 0.05) in the present study, but these did not remain significant after multiple-testing correction. In contrast, both cecal and fecal samples from females and fecal samples from males revealed no significant differences between genera after FDR correction across genotypes.

To complement these relative abundance comparisons, we also applied ALDEx2 compositional differential abundance analysis. This analysis evaluates taxa across the entire genus-level dataset using the centered log-ratio (CLR) transformation to consider the compositional structure of 16S rRNA sequencing data. Although several taxa showed variation in effect size across groups, no single genus was identified as significantly differentially abundant after FDR correction. Together, these findings indicate that gut microbial relative abundance comparisons among the top dominant genera reveal limited genotype-associated differences in FXS.

In our study, the interconnected aspect of the gut microbiota-epithelial barrier is intriguing. Our findings demonstrate molecular signatures of impaired barrier integrity in both *Fmr1* KO and Het females. qRT-PCR analyses revealed significant downregulation of core tight-junction regulators—*Tjp1, Ocln* in both KO and Het females, indicating a genotype-dependent weakening of epithelial cohesion. The pore-forming *Cldn2* was not significantly altered in females across genotypes, however the relative *Cldn2* expression was highest in Het and lowest in KO. Upregulated *Cldn2* is prominent in inflammation and immune-related gut dysfunction, leading to intestinal hyperpermeability through a leak-flux mechanism (50). In Het females, mosaic FMRP expression likely creates a mixed epithelial environment in which some cells experience greater functional stress than others (51). During embryonic stages, the cells randomly select X chromosome activation among the two X chromosomes in females (52), which may partly explain increased *Cldn2* expression in Hets. This suggests that partial, heterogeneous loss of FMRP may destabilize tight-junction regulation more than complete loss, producing elevated *Cldn2* levels observed in the Het group compared to KO females. We measured another transcriptional factor, important for assessing gut barrier integrity, *Plvap* (53). *Plvap* plays a pivotal role in maintaining endothelial homeostasis and regulating vascular permeability (54, 55). Decreased *Plvap* mRNA promotes severe protein-losing enteropathy as measured with *in vivo* studies (55). Downregulated *Plvap* may further establish uncontrollable vascular nutrient exchange addressing catastrophic plasma protein loss, despite preserved epithelial barrier integrity. Interestingly, in our study the *Plvap* expression was significantly upregulated in WT females compared to both Hets and KO females, suggesting disrupted endothelial barrier integrity in FXS.

Furthermore, female KO mice displayed reduced expression of *Muc2* and *Reg3g*, genes essential for mucosal barrier and anti-microbial defense system, respectively (56, 57). Together, these transcriptional shifts suggest that FMRP loss may predispose the gut compromised epithelium to decreased barrier stability and altered anti-microbial protection in FXS. In contrast, *Alpi* is typically reduced under conditions of dysbiosis, increasing permeability, and heightening inflammatory signaling (58). In our study, *Alpi* expression did not differ significantly across genotypes. This *Alpi* expression pattern was consistent in both male and female mice, indicating preservation of *Alpi*-associated barrier function despite *Fmr1* loss. We also examined *Tnf* which is known to indirectly disrupt tight-junction proteins in ASD and FXS through inflammatory signaling pathways (59). However, we did not detect significant alterations in *Tnf* expression either in males or females across genotype. The observed significant downregulation of *Tjp1, Ocln, Plvap, Muc2*, and *Reg3g* in FXS mice therefore likely suggests that gut barrier impairment may arise through mechanisms independent of overt *Tnf* elevation.

An unexpected yet intriguing observation was observed in female Het mice, showing a distinct gut microbiome together with elevated epithelial barrier function compared to WT females. Increased diversity is often associated with improved microbial ecosystem resilience and functional redundancy (60). Our physiological data obtained from Ussing chamber analysis revealed that the highest transepithelial resistance was in Het females. Additionally, Het females showed the highest diversity in Chao1 index in both fecal and cecal contents. This finding further suggests that a greater microbial richness may reinforce functional epithelial barrier in FXS. This observation directs us to another possibility about the mosaic *Fmr1* expression in Het females. Mosaicism produces a unique host environment that potentially enables a selective gut environment favoring a more beneficial and stable microbiota. Further investigation warrants exploration of whether this reflects an adaptive compensatory mechanism or an inherently distinct host–microbe interaction pattern in Het females. Pearce et al., (2018), showed that *Tjp*1 and *Ocln* regulate macromolecular permeability most. Even though in our experiment, we found no significant difference for *Tjp*1 and *Ocln* expression in males, in KO males, I_sc_ level was higher compared to WT, suggesting transcriptional markers may have a limited influence on transepithelial electrical properties.

In contrast, neither female nor male KO mice exhibited enhanced gut barrier function, suggesting that complete *Fmr1* loss may not solely compromise host intestinal barrier properties. As mentioned earlier, FMRP regulates the translation of a large network of target mRNAs (62). Therefore, the complete loss of *Fmr1* might induce compensatory transcriptional and post-transcriptional mechanisms in our *Fmr1* KO animals. In male mice, loss of *Fmr1* was associated with a significant increase in Isc compared with WT controls, indicating altered baseline electrogenic ion transport. Importantly, the increase in Isc without altered Vt, TER, and Gt also suggests *Fmr1* loss in males selectively affects active epithelial transport processes rather than epithelial barrier integrity and functional properties.

In contrast to the male phenotype, female KO mice exhibited genotype-dependent modulation of epithelial barrier properties without demonstrating shifts in electrogenic ion transport. Het females displayed significantly increased TER and reduced Gt relative to WT females, indicating enhanced epithelial barrier tightness. KO females showed a similar trend toward increased TER, although this did not reach statistical significance, and no genotype-dependent differences in Isc or Vt were detected in females across genotypes. Together, these findings indicate that *Fmr1* mutation influences epithelial electrical barrier properties in females while leaving baseline ion transport largely unaffected. Het females exhibited the highest *Cldn2* expression across genotypes, although this difference did not reach statistical significance. Important to mention that, *Cldn2* functions as a pore-forming tight junction protein typically associated with increased paracellular permeability (17, 63). Therefore, elevated *Cldn2* expression would generally be expected to reduce epithelial barrier resistance. However, Het females displayed significantly higher TER compared with WT females, suggesting that overall epithelial barrier integrity may be preserved or reinforced in Hets despite increased *Cldn2* expression.

Our findings further support the concept that transcriptional changes occur within interconnected regulatory networks rather than acting in isolation (64). In male mice, significant alterations in epithelial ion transport were observed across genotype despite the absence of detectable changes in mRNA expression related to gut barrier integrity, mucosal defense, or inflammation. This dissociation suggests that steady-state mRNA levels are not sufficient to predict epithelial function. Such functional changes may arise from post-transcriptional regulation (65, 66), including altered protein abundance, impaired trafficking, or defects in post-translational modification. Together, these findings indicate that preserved transcript levels do not necessarily guarantee effective protein function to maintain functional epithelial barrier homeostasis. Additionally, increased microbial species richness (i.e., Chao1 index) in female Het mice with elevated transepithelial barrier resistance may indicate that higher microbial diversity may be positively linked to barrier resistance. One of the justifying mechanisms behind this claim could be that higher diversity introduces more beneficial metabolites that can further act as a protective anchor for gut barrier integrity and function (67–69).

In females, genotype-dependent reductions in *Tjp1, Ocln*, and *Plvap* indicate transcriptional remodeling of barrier components. Yet these changes did not uniformly translate into barrier dysfunction. Together, these data highlight that mRNA changes may have a limited functional impact, while physiological alterations can arise independently of major transcriptional shifts. Collectively, this study identifies sex- and genotype-specific perturbations in gut microbial landscape and epithelial barrier characteristics associated with FXS. The overall findings underscore the importance of considering sex chromosomes and mosaicism when evaluating gut phenotypes in X-linked neurodevelopmental disorders, like FXS. Nevertheless, our study also has some limitations. For example, differences in microbiome and barrier outcomes may be influenced by X-chromosome mosaicism in Hets, complicating the interpretation of heterozygous phenotypes. Furthermore, regional sampling constraints in the intestine may not fully capture spatial heterogeneity in epithelial physiology or integrity (70). 16S rRNA sequencing provides taxonomic resolution but limited functional insight, preventing direct identification of microbial metabolic pathways driving barrier differences. Meanwhile, Ussing chamber measurements, while precise, reflect *ex vivo* physiology and may not capture dynamic immune–microbe–epithelium interactions occurring *in vivo*. Yet, our study also has interesting findings. Integrating microbiome profiling with barrier physiology, this study provides a comprehensive systems-level view of GI phenotypes in FXS. Inclusion of both sexes and heterozygous females enables detection of sex-linked and X-chromosome–dependent effects relevant to human FXS. Moreover, using tissue-specific physiological assays allows us to directly assess epithelial properties that cannot be measured non-invasively in humans. Future work integrating metagenomics, metabolomics, and cellular analyses will be worth investigating to identify specific microbial taxa, metabolites, and epithelial signaling pathways that mediate these effects in FXS. Understanding these interactions may ultimately support the development of microbiome- or barrier-targeted interventions for tackling GI symptoms frequently experienced by FXS patients.

Collectively, our findings demonstrate that loss of FMRP is associated with alterations in gut microbiome composition alongside changes in intestinal epithelial barrier regulation in mice. While complete *Fmr1* loss did not uniformly impair gut barrier integrity, distinct physiological and microbial signatures were observed across genotypes in females, particularly in female heterozygous mice. In summary, these results highlight a previously underappreciated link between FMRP deficiency, host–microbiome interactions, and epithelial barrier regulation, providing new insights into GI alterations in FXS.

## Data Availability

The 16S rRNA gene sequencing data generated in this study have been deposited in the NCBI Sequence Read Archive (SRA) under BioProject accession number PRJNA1429409. These data will be publicly available upon release on August 11, 2027. Until that date, the data remain archived in the SRA under temporary submission ID SUB15983690.

## Notes

### Competing Interest Statement

The authors have declared no competing interest.

### Funding

- NIH Common Fund: OCMR COBRE

