## supplemental figures and tables for "*Fmr1* mutation reshapes gut microbiome structure, diversity, intestinal barrier integrity, and function in a sex and genotype-dependent manner"

| Gene | Forward | Reverse |
| --- | --- | --- |
| *mReg3g* | 5`-cca tct tca cgt agc agc-3` | 5`-caa gat gtc ctg agg gc-3` |
| *mTnf* | 5`-ctg agg tca atc tgc cca agt ac-3` | 5`-ctt cac aga gca atg act cca aag-3` |
| *mMuc2* | 5`-ctg acc aag agc gaa cac aa-3` | 5`-cat gac tgg aag caa ctg ga-3` |
| *mcldn2* | 5`-cgg ctc cgt ttt cta gat gc-3` | 5`-ggc tgc tgc tct tgc ttc tt -3` |
| *mplvap* | 5`- gcc agg tgg ttg gac tat ctg-3` | 5`- ctc cat ctc acg tcg cgt a-3` |
| *mOcln* | 5`-acc cga aga aag atg gat tcg -3` | 5`-cat agt cag atg ggg gtg ga -3` |
| *mTjp1* | 5`-agc ctg gtt gtt tag gag ca -3` | 5`- cag aat acg gct cct tcc tg-3` |
| *mAlpi* | 5`-agg aca tcg cca ctc aac tc -3` | 5`-ggt tcc aga ctg gtt act gtc a -3` |
| *mGapdh* | 5`-caa ggt cat cca tga caa ctt tg -3` | 5`- ggc cat cca cag tct tct gg-3` |

**Extended Table 1 | Primer sequences for gene expression analysis**

**Extended Table 2 | Genus-Level Relative Abundance Differences Between cecal WT and KO males**

| **Genus** | **WT_mean** | **KO_mean** | **p_value** | **FDR** |
| --- | --- | --- | --- | --- |
| *Clostridium sensu stricto 1* | 4.36E-06 | 0.000482 | 0.060808 | 0.133777 |
| *Lachnoclostridium* | 0.040563 | 0.028953 | 0.008658 | 0.047619 |
| Lachnospiraceae_NK4A136_group | 0.029681 | 0.039033 | 0.393939 | 0.481481 |
| Lachnospiraceae_UCG_006 | 0.025359 | 0.037622 | 0.24026 | 0.377551 |
| *Lactobacillus* | 0.020342 | 0.00268 | 0.004329 | 0.047619 |
| *Muribaculum* | 0.081753 | 0.051841 | 0.025974 | 0.071429 |
| *Turicibacter* | 0.026639 | 0.095053 | 0.015152 | 0.055556 |
| unclassified_Lachnospiraceae | 0.073727 | 0.050934 | 0.132035 | 0.242063 |
| unclassified_Muribaculaceae | 0.36825 | 0.374375 | 0.937229 | 0.937229 |

**9**

**Extended Table 3 | Genus-Level Relative Abundance Differences Between cecal WT, KO and Het females**

| Genus | WT_mean | KO_mean | Het_mean | p_value | FDR |
| --- | --- | --- | --- | --- | --- |
| *Clostridium sensu stricto_1* | 0.020477 | 0.046904 | 0.009573 | 0.196597 | 0.480444 |
| *Lachnoclostridium* | 0.027136 | 0.024887 | 0.043548 | 0.046413 | 0.480444 |
| Lachnospiraceae_NK4A136_group | 0.064639 | 0.035122 | 0.028368 | 0.219892 | 0.480444 |
| Lachnospiraceae_UCG_006 | 0.01881 | 0.046712 | 0.035135 | 0.26206 | 0.480444 |
| *Muribaculum* | 0.036139 | 0.039575 | 0.050894 | 0.240052 | 0.480444 |
| *Lactobacillus* | 0.062325 | 0.10938 | 0.115085 | 0.323468 | 0.508307 |
| *Turicibacter* | 0.050279 | 0.105866 | 0.086684 | 0.475832 | 0.654269 |
| unclassified_Lachnospiraceae | 0.054608 | 0.050035 | 0.059361 | 0.75085 | 0.830779 |
| unclassified_Muribaculaceae | 0.308029 | 0.286385 | 0.274444 | 0.755254 | 0.830779 |

**Extended Table 4 | Genus-Level Relative Abundance Differences Between fecal WT and KO males**

| **Genus** | **WT_mean** | **KO_mean** | **p_value** | **FDR** |
| --- | --- | --- | --- | --- |
| *Clostridium sensu stricto 1* | 9.86E-05 | 0.009954 | 0.326058 | 0.92517 |
| Lachnospiraceae_NK4A136_group | 0.038132 | 0.03288 | 0.588745 | 0.92517 |
| *Lactobacillus* | 0.10714 | 0.109062 | 0.588745 | 0.92517 |
| *Muribaculum* | 0.022612 | 0.019074 | 0.132035 | 0.92517 |
| unclassified_Muribaculaceae | 0.211133 | 0.259257 | 0.393939 | 0.92517 |
| *Turicibacter* | 0.08792 | 0.074134 | 0.699134 | 0.96131 |
| *Lachnoclostridium* | 0.021636 | 0.023777 | 1 | 1 |
| Lachnospiraceae_UCG_006 | 0.00681 | 0.03023 | 0.937229 | 1 |
| unclassified_Lachnospiraceae | 0.02964 | 0.032508 | 0.937229 | 1 |

**Extended Table 5 | Genus-Level Relative Abundance Differences among fecal WT, KO and Het females**

| **Genus** | **WT_mean** | **KO_mean** | **Het_mean** | **p_value** | **FDR** |
| --- | --- | --- | --- | --- | --- |
| *Turicibacter* | 0.13718 | 0.151499 | 0.044241 | 0.007844 | 0.086289 |
| *Clostridium sensu stricto 1* | 0.058773 | 0.084982 | 0.00606 | 0.039271 | 0.215993 |
| *Muribaculum* | 0.036385 | 0.037277 | 0.020571 | 0.140168 | 0.385462 |
| unclassified_Muribaculaceae | 0.247606 | 0.298005 | 0.226139 | 0.139351 | 0.385462 |
| unclassified_Lachnospiraceae | 0.028679 | 0.019673 | 0.028507 | 0.367879 | 0.674446 |
| *Lactobacillus* | 0.200819 | 0.154652 | 0.256667 | 0.484254 | 0.76097 |
| Lachnospiraceae_NK4A136_group | 0.03575 | 0.023375 | 0.020211 | 0.587333 | 0.807583 |
| *Lachnoclostridium* | 0.016342 | 0.018547 | 0.027801 | 0.884434 | 0.926795 |
| Lachnospiraceae_UCG_006 | 0.010802 | 0.016516 | 0.040568 | 0.926795 | 0.926795 |

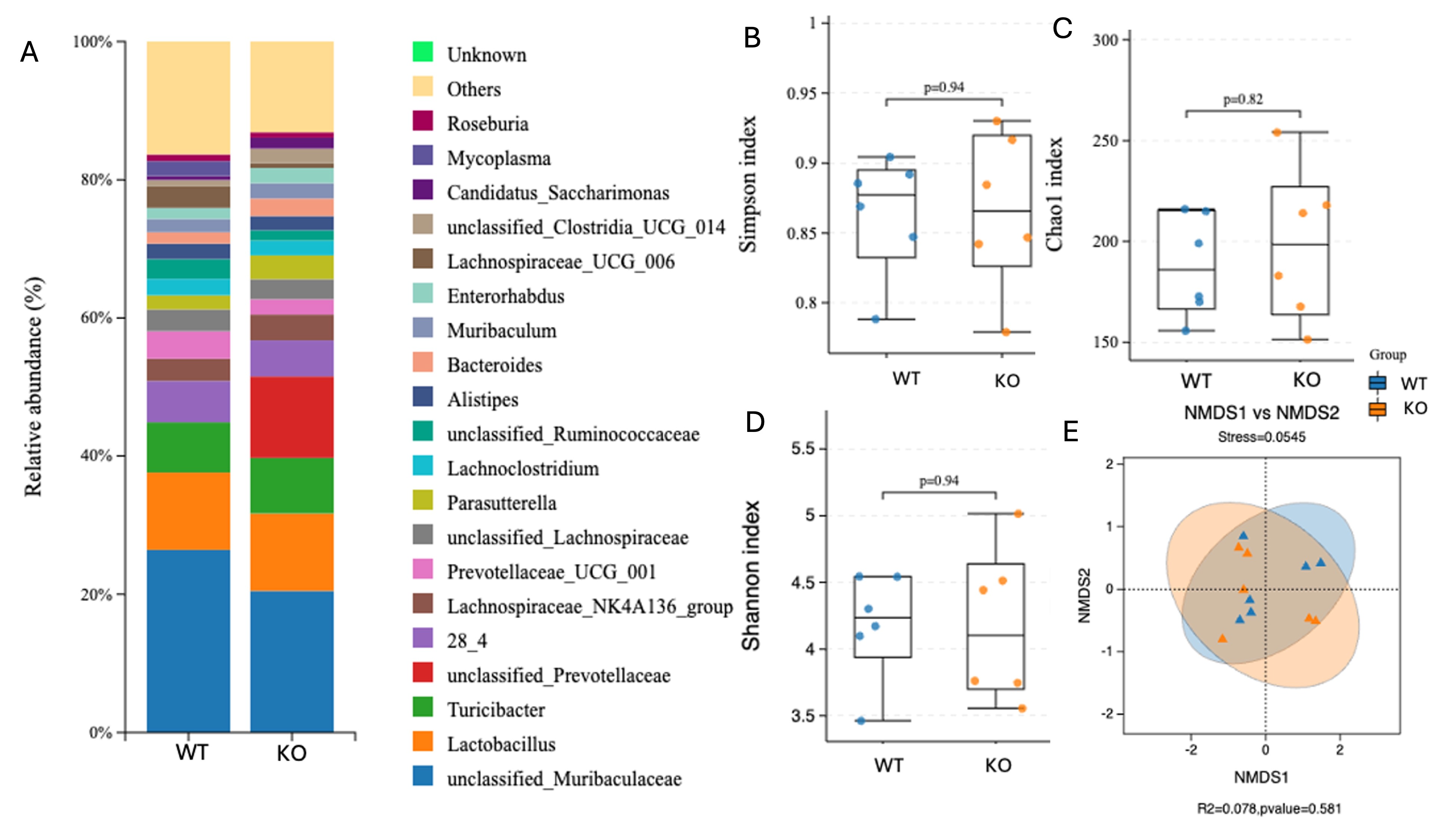
 **Supplemental Figure 1 | Fecal microbiome composition and microbial diversity in male WT and KO mice.**

Fecal microbial communities from WT and KO male mice were profiled by 16S rRNA gene sequencing. (A) Relative abundance of the top 20 bacterial genera in fecal contents from WT and KO male mice. Genus-level microbial composition displayed as stacked bar plots showing relative abundance.; low-abundance taxa are grouped as “Others,” and unassigned taxa are denoted as “Unknown.” (B–D) Alpha-diversity metrics comparing WT and KO males, including the Simpson index (B), which reflects community evenness and dominance, the Chao1 index (C), an estimator of species richness, and the Shannon diversity index (D), which integrates both richness and evenness. Statistical significance was assessed using the Mann–Whitney U test. Each point represents an individual animal, and box plots depict the median and interquartile range.
(E) Beta-diversity analysis based on Bray–Curtis dissimilarity at the species level, visualized by non-metric multidimensional scaling (NMDS). Group differences were evaluated using PERMANOVA, with shaded ellipses representing 95% confidence intervals for each genotype.
All microbiome bioinformatic analyses were performed using the BMKCloud platform ([www.biocloud.net](http://www.biocloud.net)). Data are derived from six independent biological replicates per group (n = 6 per genotype).

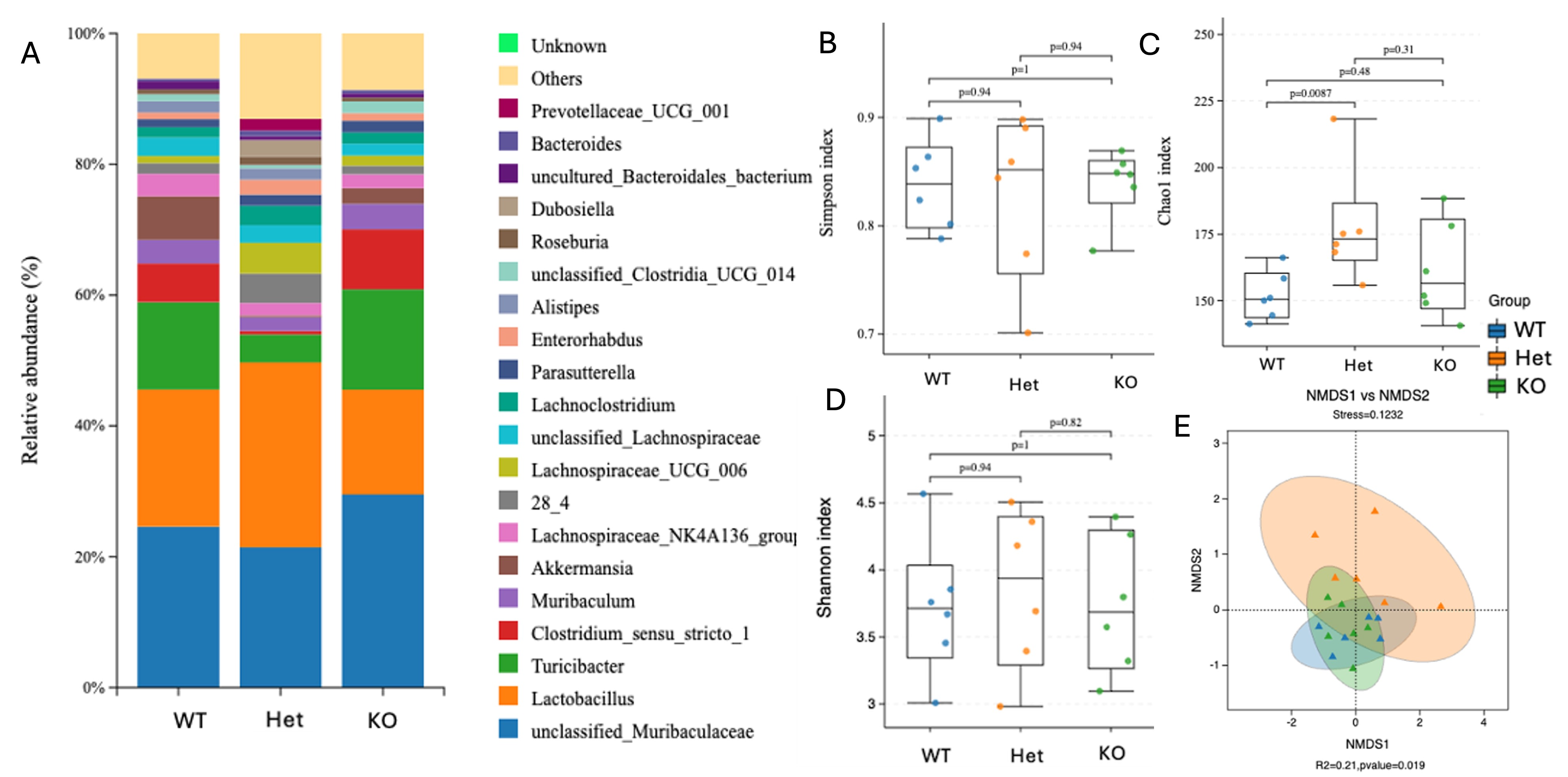
**Supplemental Figure 2 | Fecal microbiome composition and microbial diversity among female WT, Het and KO mice.**

Fecal microbial communities from WT, Het and KO female mice were profiled by 16S rRNA gene sequencing. (A) Relative abundance of the top 20 bacterial genera in fecal contents among WT, Het and KO female mice. Genus-level microbial composition displayed as stacked bar plots showing relative abundance.; low-abundance taxa are grouped as “Others,” and unassigned taxa are denoted as “Unknown.” (B–D) Alpha-diversity metrics comparing WT and KO males, including the Simpson index (B), which reflects community evenness and dominance, the Chao1 index (C), an estimator of species richness, and the Shannon diversity index (D), which integrates both richness and evenness. Statistical significance was assessed using the Mann–Whitney U test. Each point represents an individual animal, and box plots depict the median and interquartile range. (E) Beta-diversity analysis based on Bray–Curtis dissimilarity at the species level, visualized by non-metric multidimensional scaling (NMDS). Group differences were evaluated using PERMANOVA, with shaded ellipses representing 95% confidence intervals for each genotype. All microbiome bioinformatic analyses were performed using the BMKCloud platform. Data are derived from six independent biological replicates per group (n = 6 per genotype).

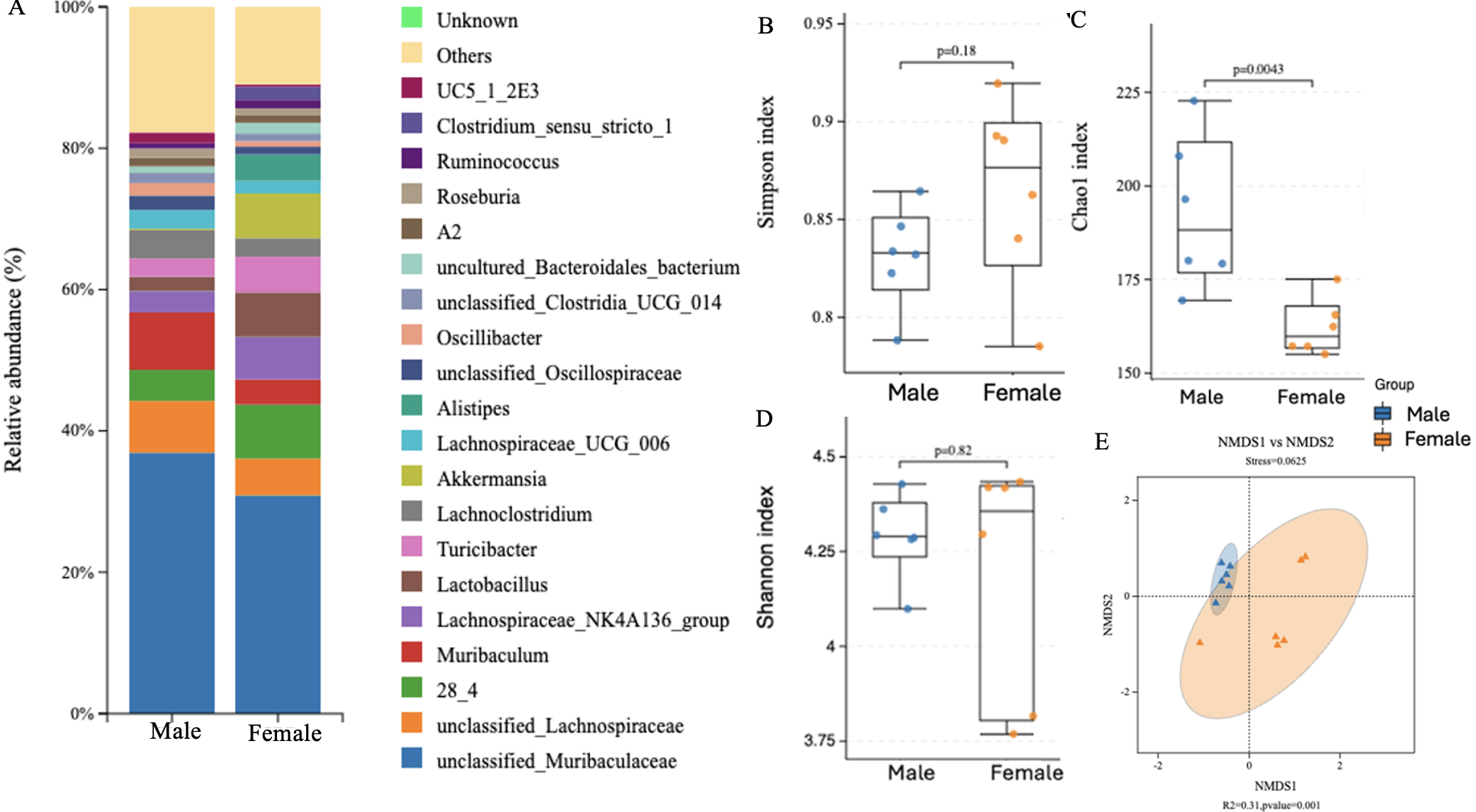
 **Supplemental Figure 3 | Sex-associated differences in cecal microbiome composition and microbial diversity in WT male and WT female mice.**

Cecal microbial communities from WT male and WT female mice were profiled by 16S rRNA gene sequencing. (A) Relative abundance of the top 20 bacterial genera in cecal contents from WT male and WT female mice. Genus-level microbial composition displayed as stacked bar plots showing relative abundance.; low-abundance taxa are grouped as “Others,” and unassigned taxa are denoted as “Unknown.” (B–D) Alpha-diversity metrics comparing WT and KO males, including the Simpson index (B), which reflects community evenness and dominance, the Chao1 index (C), an estimator of species richness, and the Shannon diversity index (D),which integrates both richness and evenness, and ,. Statistical significance was assessed using the Mann–Whitney U test. Each point represents an individual animal, and box plots depict the median and interquartile range. (E) Beta-diversity analysis based on Bray–Curtis dissimilarity at the species level, visualized by non-metric multidimensional scaling (NMDS). Group differences were evaluated using PERMANOVA, with shaded ellipses representing 95% confidence intervals for each genotype. All microbiome bioinformatic analyses were performed using the BMKCloud platform. Data are derived from six independent biological replicates per group (n = 6 per genotype).

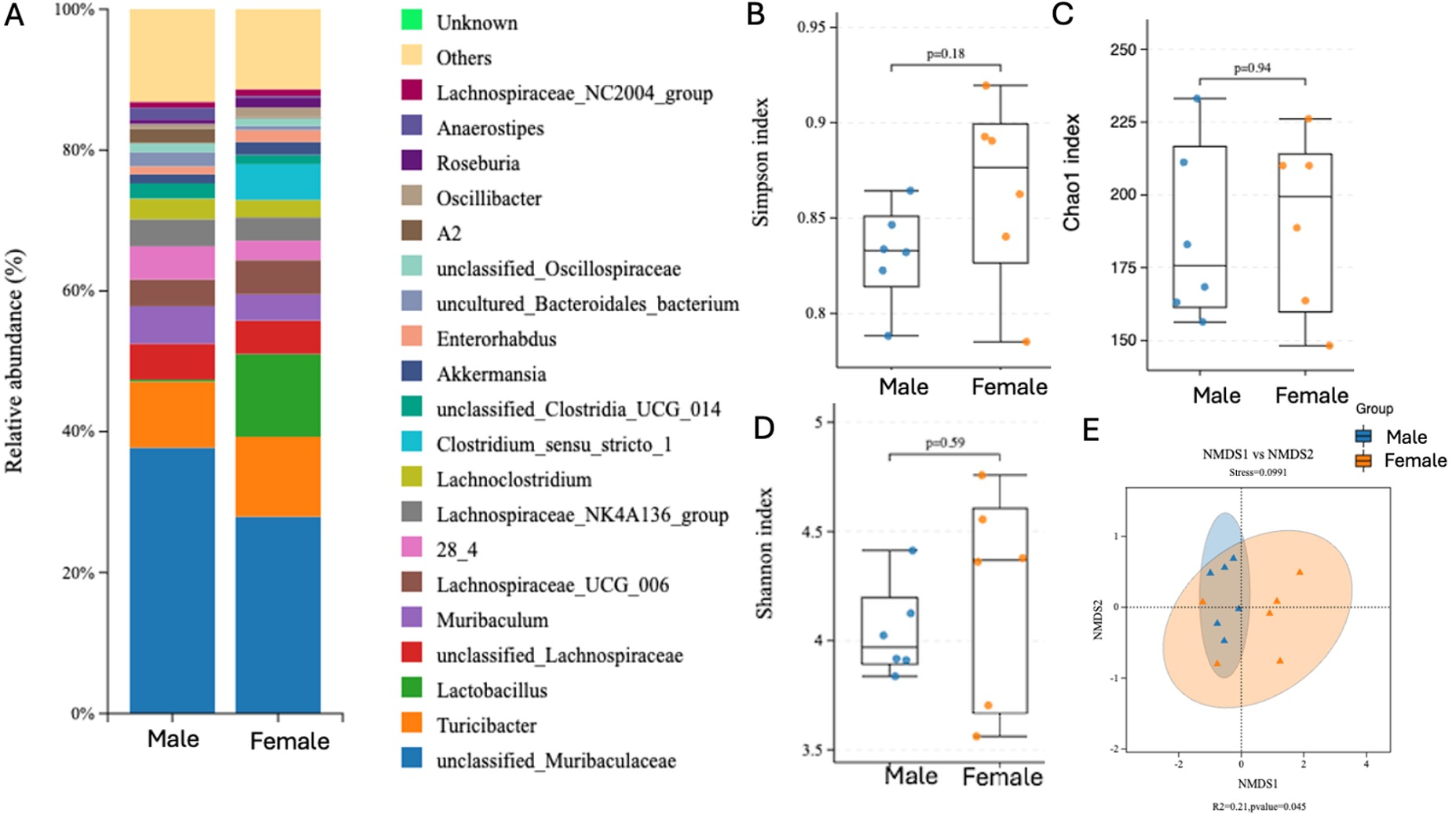
 **Supplemental Figure 4 | Sex-associated differences in cecal microbiota composition and diversity in KO male and KO female mice.**

Cecal microbial communities from KO male and KO female mice were profiled by 16S rRNA gene sequencing. (A) Relative abundance of the top 20 bacterial genera in cecal contents from KO male and KO female mice. Genus-level microbial composition displayed as stacked bar plots showing relative abundance.; low-abundance taxa are grouped as “Others,” and unassigned taxa are denoted as “Unknown.” (B–D) Alpha-diversity metrics comparing WT and KO males, including the Simpson index (B), which reflects community evenness and dominance, the Chao1 index (C), an estimator of species richness, and the Shannon diversity index (D), which integrates both richness and evenness. Statistical significance was assessed using the Mann–Whitney U test. Each point represents an individual animal, and box plots depict the median and interquartile range. (E) Beta-diversity analysis based on Bray–Curtis dissimilarity at the species level, visualized by non-metric multidimensional scaling (NMDS). Group differences were evaluated using PERMANOVA, with shaded ellipses representing 95% confidence intervals for each genotype. All microbiome bioinformatic analyses were performed using the BMKCloud platform. Data are derived from six independent biological replicates per group (n = 6 per genotype).

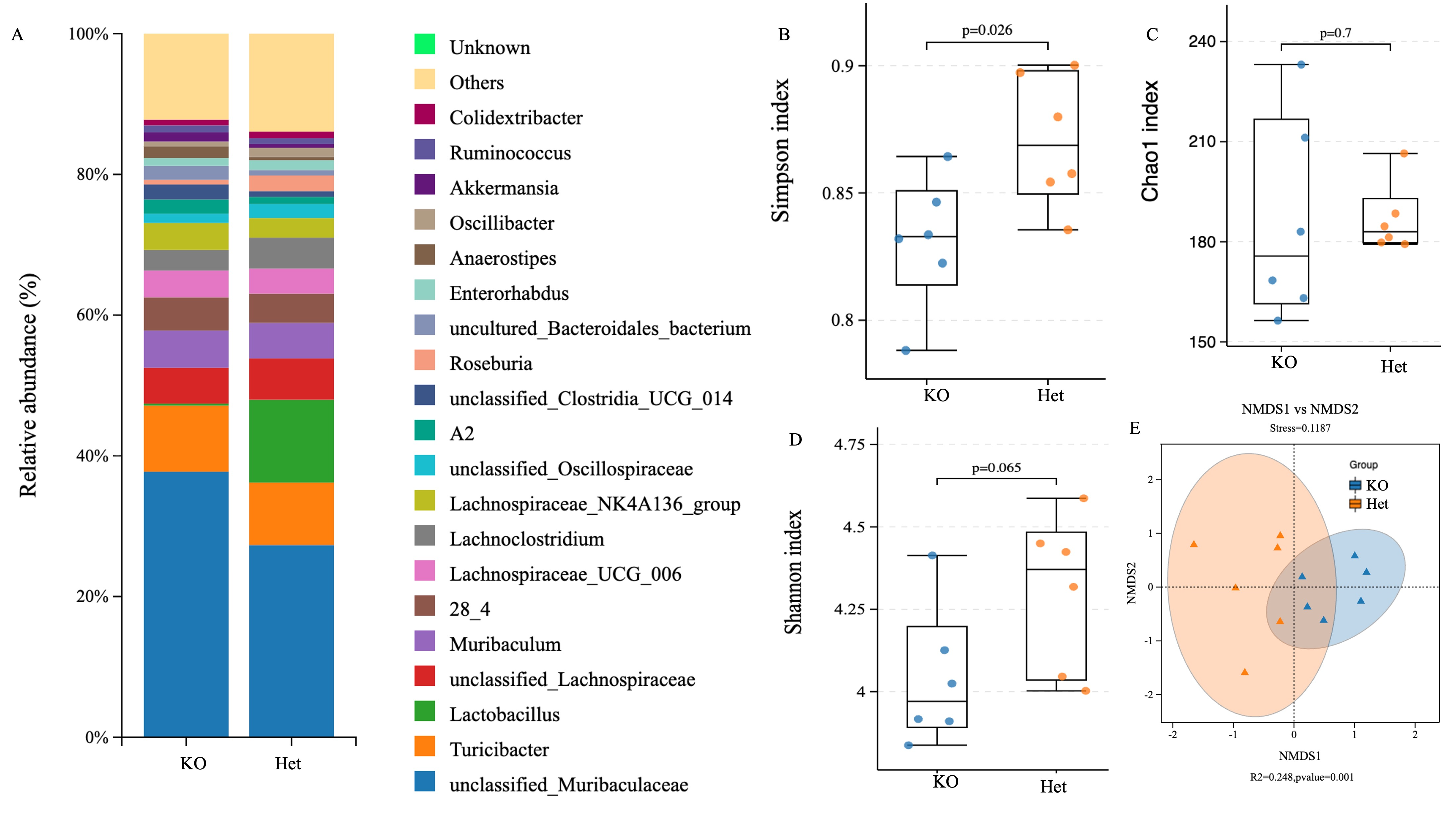

**Supplemental Figure 5 | Gut microbial community structure and diversity in KO males and Heterozygous (Het) females in cecal contents**

Cecal microbial communities and their diversity from KO male and Het female mice were profiled by 16S rRNA gene sequencing. (A) Relative abundance of the top 20 bacterial genera in fecal contents from KO male and Het female mice. Genus-level microbial composition displayed as stacked bar plots showing relative abundance.; low-abundance taxa are grouped as “Others,” and unassigned taxa are denoted as “Unknown.” (B–D) α-diversity metrics, including Simpson index (B), Shannon diversity index (C), and Chao1 richness (D), presented as box-and-whisker plots depicting the median, interquartile range, and full data range, with individual points representing independent biological replicates. Statistical comparisons for α-diversity were performed using the Mann–Whitney U test. (E) β-diversity analysis visualized by non-metric multidimensional scaling (NMDS) based on Bray–Curtis dissimilarity at the species level, with 95% confidence ellipses indicating group dispersion. Group differences in community structure were assessed using PERMANOVA. Bioinformatic processing and visualization for panels A–E were performed using the BMKCloud platform. Each group included six independent biological replicates (n = 6 per group). Exact P values, stress values, and R² statistics are indicated in the respective panels.

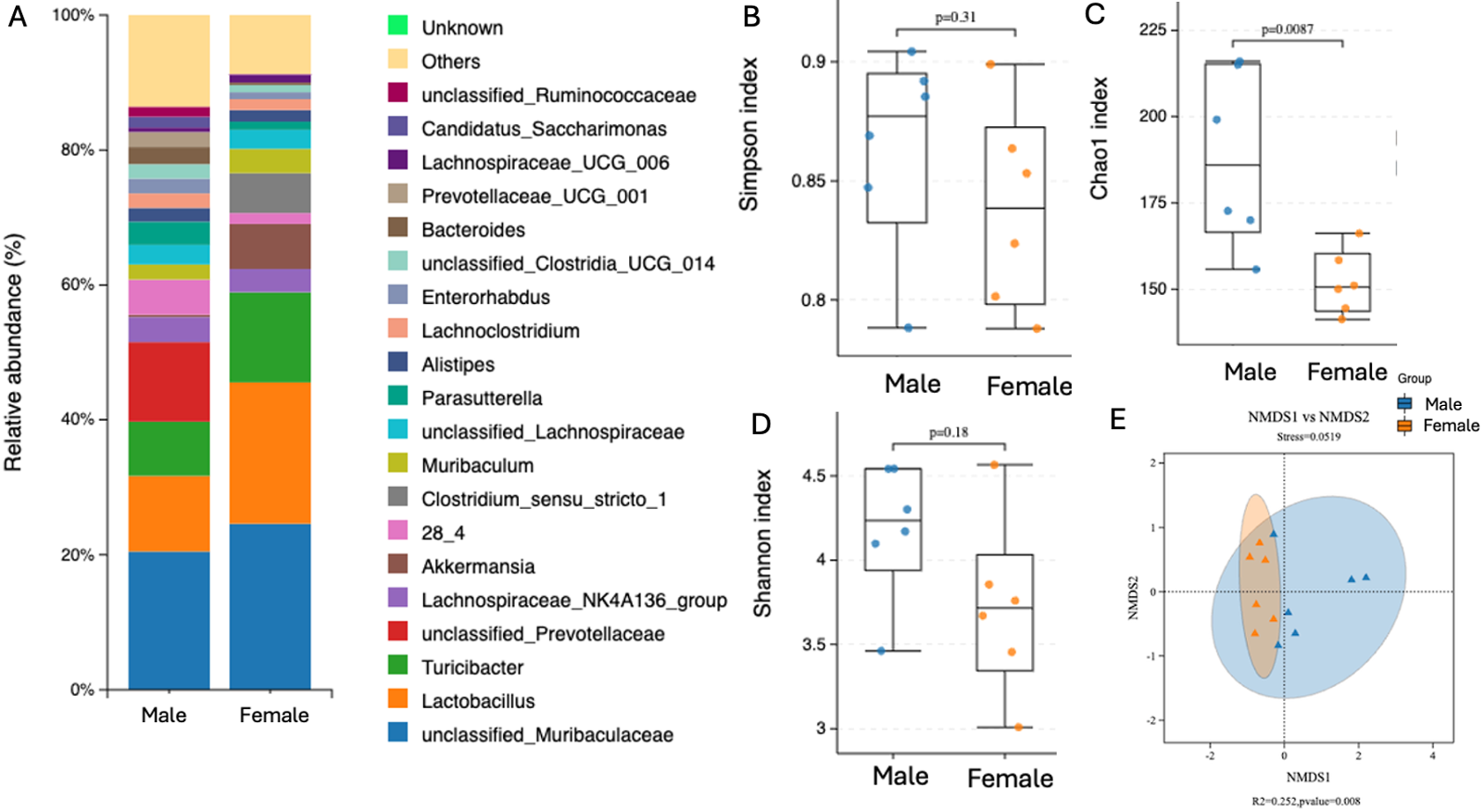

**Supplemental Figure 6 | Sex-associated differences in fecal microbiome composition and microbial diversity in WT male and WT female mice.**

Fecal microbial communities from WT male and WT female mice were profiled by 16S rRNA gene sequencing. (A) Relative abundance of the top 20 bacterial genera in fecal contents from WT male and WT female mice. Genus-level microbial composition displayed as stacked bar plots showing relative abundance.; low-abundance taxa are grouped as “Others,” and unassigned taxa are denoted as “Unknown.” (B–D) Alpha-diversity metrics comparing WT and KO males, including the Simpson index (B), which reflects community evenness and dominance, the Chao1 index (C), an estimator of species richness, and the Shannon diversity index (D), which integrates both richness and evenness. Statistical significance was assessed using the Mann–Whitney U test. Each point represents an individual animal, and box plots depict the median and interquartile range. (E) Beta-diversity analysis based on Bray–Curtis dissimilarity at the species level, visualized by non-metric multidimensional scaling (NMDS). Group differences were evaluated using PERMANOVA, with shaded ellipses representing 95% confidence intervals for each genotype. All microbiome bioinformatic analyses were performed using the BMKCloud platform. Data are derived from six independent biological replicates per group (n = 6 per genotype).

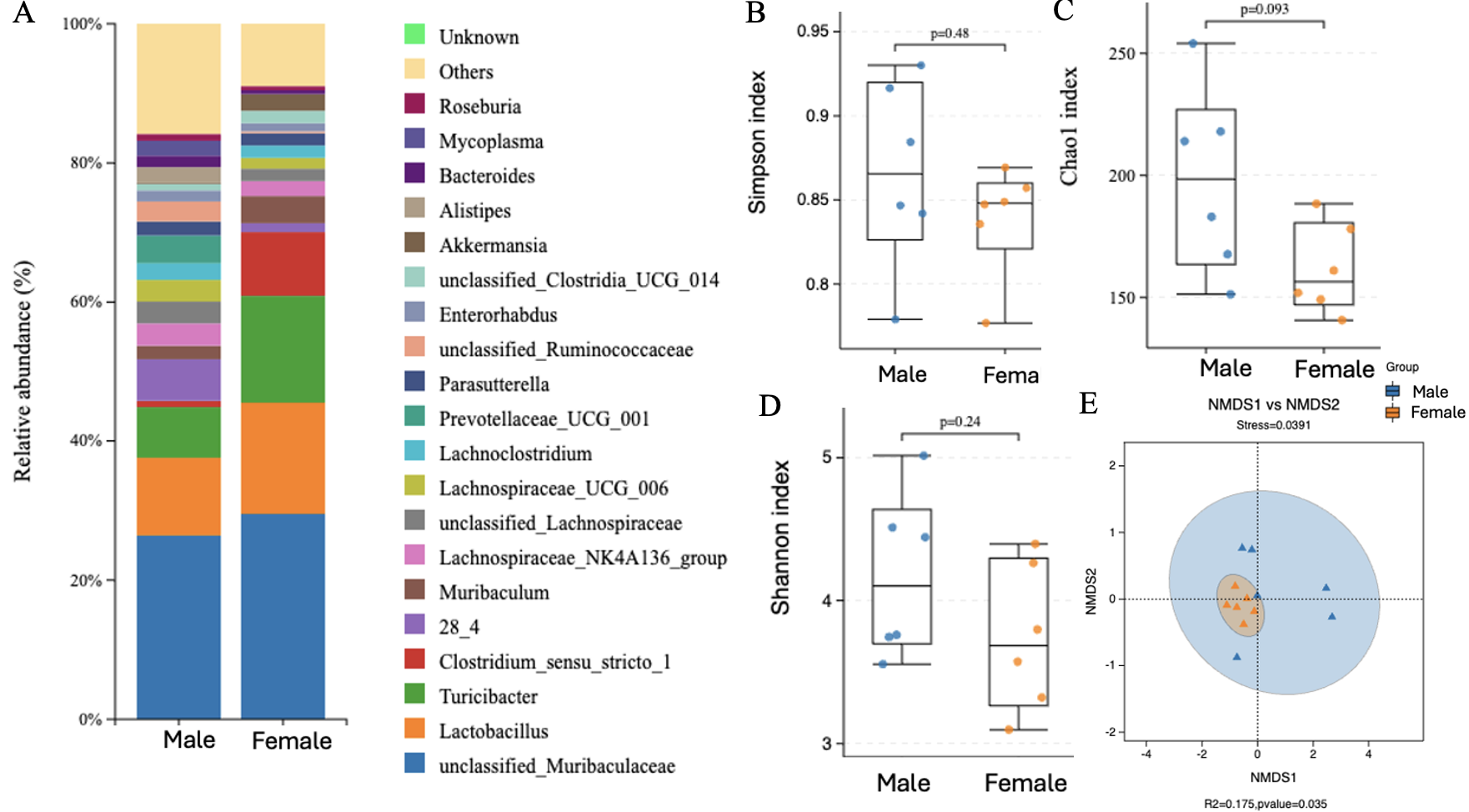
 **Supplemental Figure 7 | Sex-associated differences in fecal microbiome composition and microbial diversity in KO male and KO female mice.**

Fecal microbial communities from KO male and KO female mice were profiled by 16S rRNA gene sequencing. (A) Relative abundance of the top 20 bacterial genera in fecal contents from KO male and KO female mice. Genus-level microbial composition displayed as stacked bar plots showing relative abundance.; low-abundance taxa are grouped as “Others,” and unassigned taxa are denoted as “Unknown.” (B–D) α-diversity metrics, including Simpson index (B), Shannon diversity index (C), and Chao1 richness (D), presented as box-and-whisker plots depicting the median, interquartile range, and full data range, with individual points representing independent biological replicates. Statistical comparisons for α-diversity were performed using the Mann–Whitney U test. (E) β-diversity analysis visualized by non-metric multidimensional scaling (NMDS) based on Bray–Curtis dissimilarity at the species level, with 95% confidence ellipses indicating group dispersion. Group differences in community structure were assessed using PERMANOVA. Bioinformatic processing and visualization for panels A–E were performed using the BMKCloud platform. Each group included six independent biological replicates (n = 6 per group). Exact P values, stress values, and R² statistics are indicated in the respective panels.

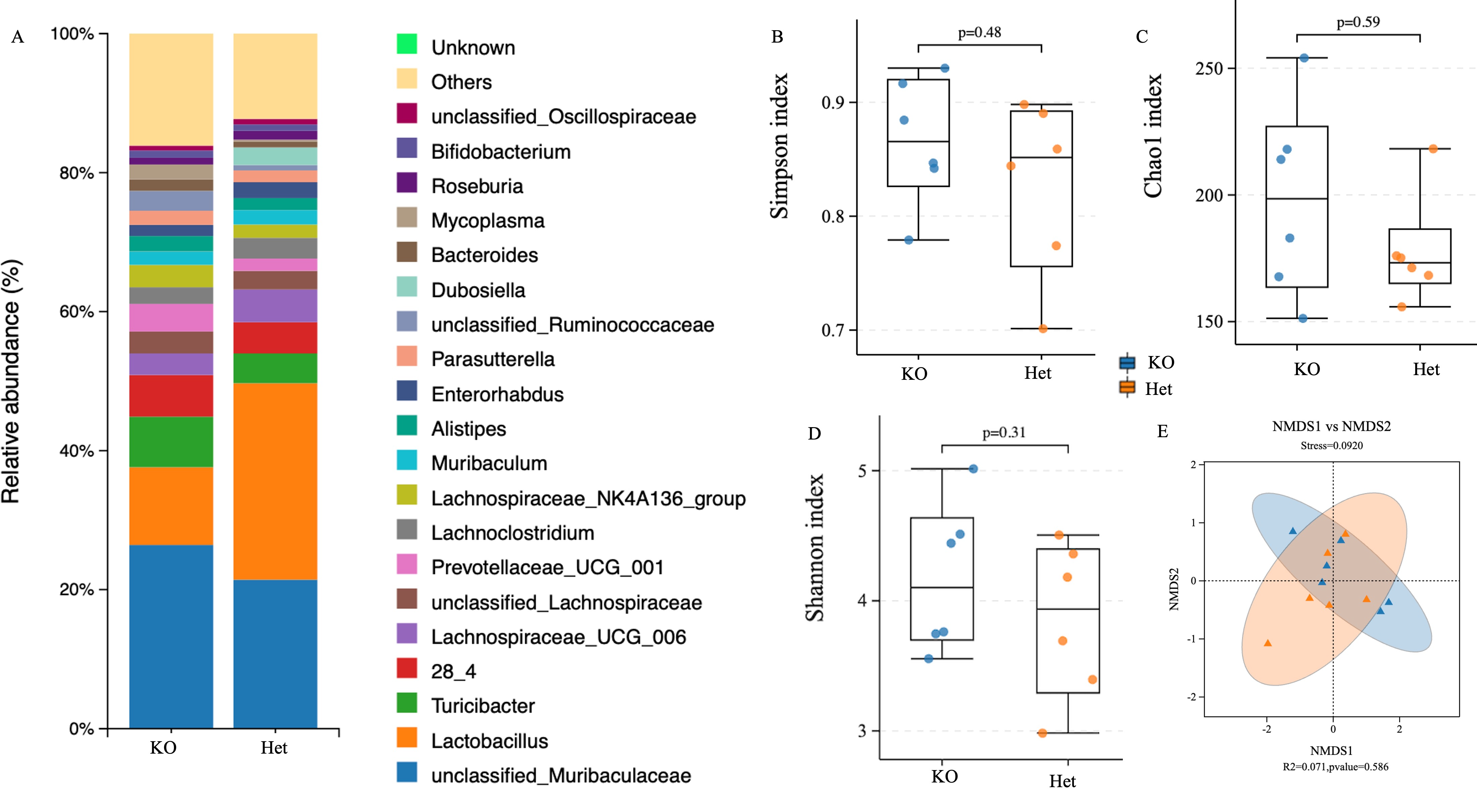

**Supplemental Figure 8 | Gut microbial community structure and diversity in KO males and Heterozygous (Het) females in fecal contents**

Fecal microbial communities and their diversity from KO male and Het female mice were profiled by 16S rRNA gene sequencing. (A) Relative abundance of the top 20 bacterial genera in fecal contents from KO male and Het female mice. Genus-level microbial composition displayed as stacked bar plots showing relative abundance.; low-abundance taxa are grouped as “Others,” and unassigned taxa are denoted as “Unknown.” (B–D) α-diversity metrics, including Simpson index (B), Shannon diversity index (C), and Chao1 richness (D), presented as box-and-whisker plots depicting the median, interquartile range, and full data range, with individual points representing independent biological replicates. Statistical comparisons for α-diversity were performed using the Mann–Whitney U test. (E) β-diversity analysis visualized by non-metric multidimensional scaling (NMDS) based on Bray–Curtis dissimilarity at the species level, with 95% confidence ellipses indicating group dispersion. Group differences in community structure were assessed using PERMANOVA. Bioinformatic processing and visualization for panels A–E were performed using the BMKCloud platform. Each group included six independent biological replicates (n = 6 per group). Exact P values, stress values, and R² statistics are indicated in the respective panels.

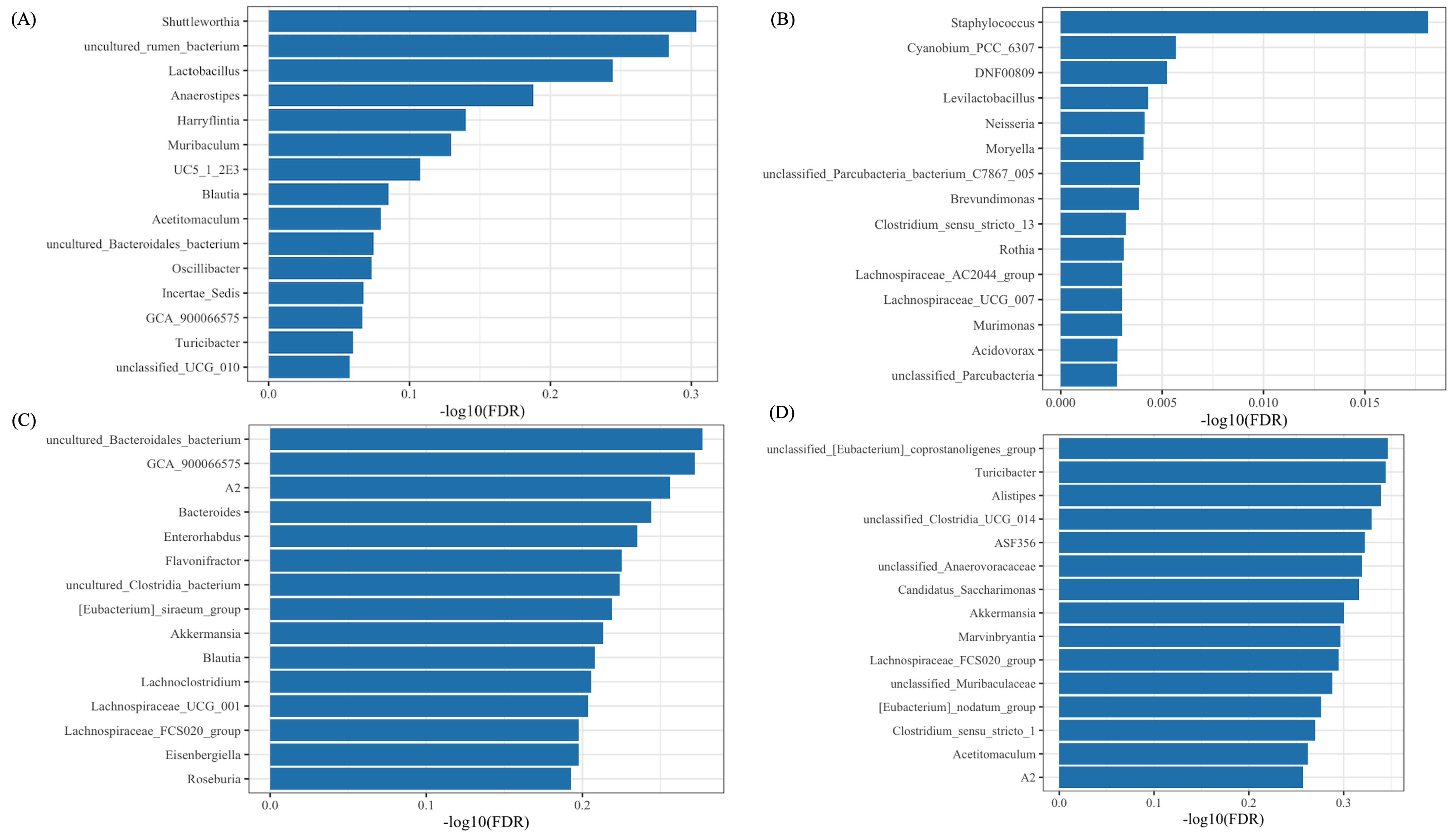

**Supplemental Figure 9 | Differential abundance of bacterial genera identified by ALDEx2**

A) Differentially abundant genera between wild-type (WT) and knockout (KO) male mice in cecal samples.
(B) Differentially abundant genera between WT and KO male mice in fecal samples.
(C) Differentially abundant genera among WT, Heterozygous (Het), and KO female mice in cecal samples.
(D) Differentially abundant genera among WT, Het, and KO female mice in fecal samples. Differential abundance was determined using ALDEx2 with Benjamini–Hochberg false discovery rate (FDR) correction. The x-axis represents −log10(FDR), and values greater than 1.3 correspond to statistically significant differences (FDR < 0.05)

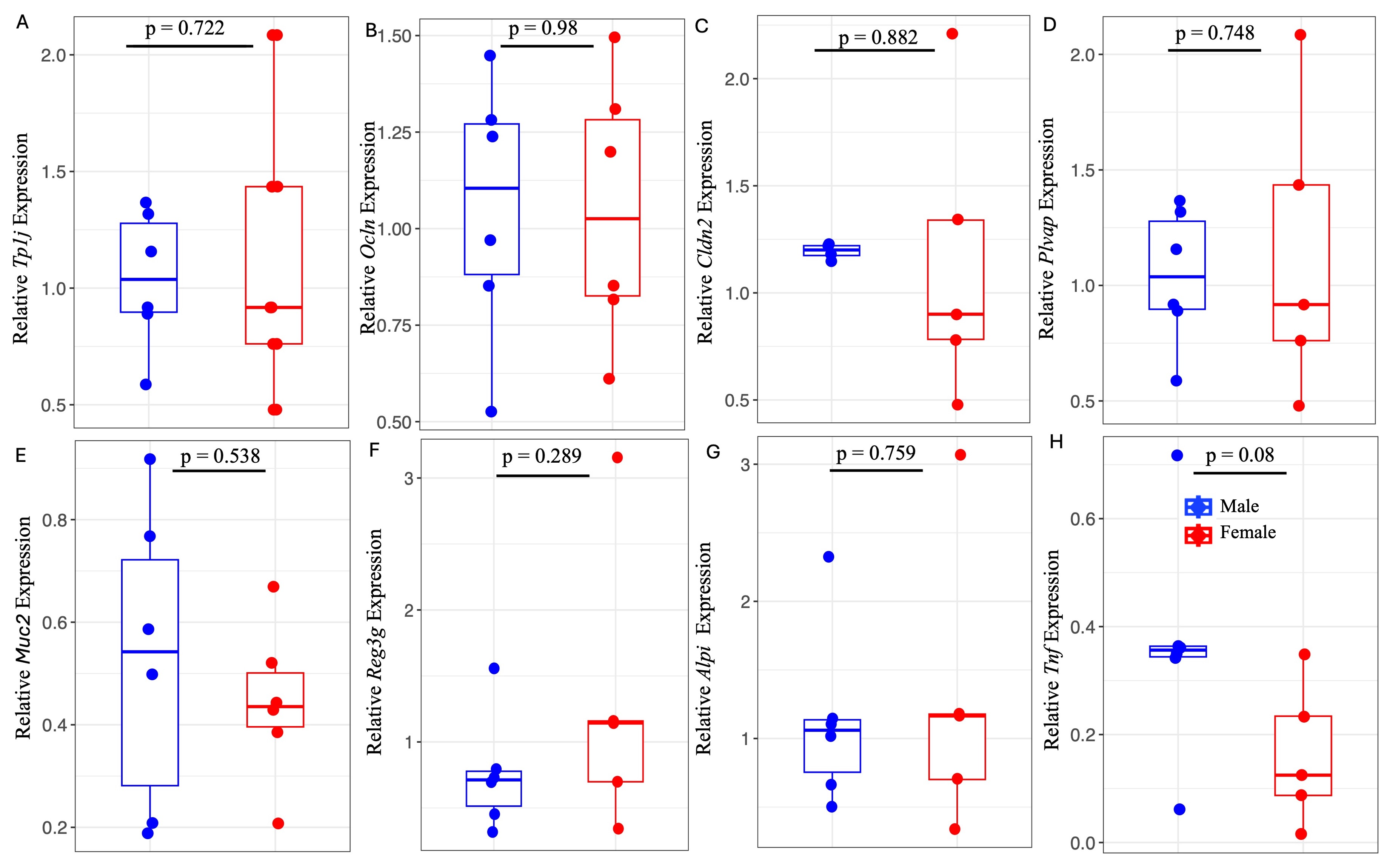

**Supplemental Figure 10 | Transcriptional markers of intestinal barrier structure, mucosal defense, and inflammatory tone across WT males and females**

Relative expression of genes spanning multiple functional layers of the intestinal barrier in WT male and WT female mice. (A) *Tjp1* and (B) *Ocln*, core tight junction–associated proteins; (C) *cldn2*, a pore-forming tight junction component; (D) *plvap,* an endothelial permeability marker; (E) *Muc2*, a mucus layer structural component; (F) *Reg3g*, an epithelial antimicrobial peptide; (G) *Alpi*, an anti-inflammatory epithelial enzyme; and (H) *Tnf*, a pro-inflammatory cytokine. Box-and-whisker plots depict the median (center line), interquartile range (box), and full data range (whiskers), with individual points representing independent biological replicates. Blue points/boxes represent WT males, and red points/boxes represent WT females. Statistical analyses were selected based on Shapiro–Wilk normality and homogeneity of variance testing: unpaired two-tailed Student’s t tests were applied for panels A-H. Each group included 5–7 independent biological replicates (n = 5–7 per group). Exact P values are indicated in the respective panels.

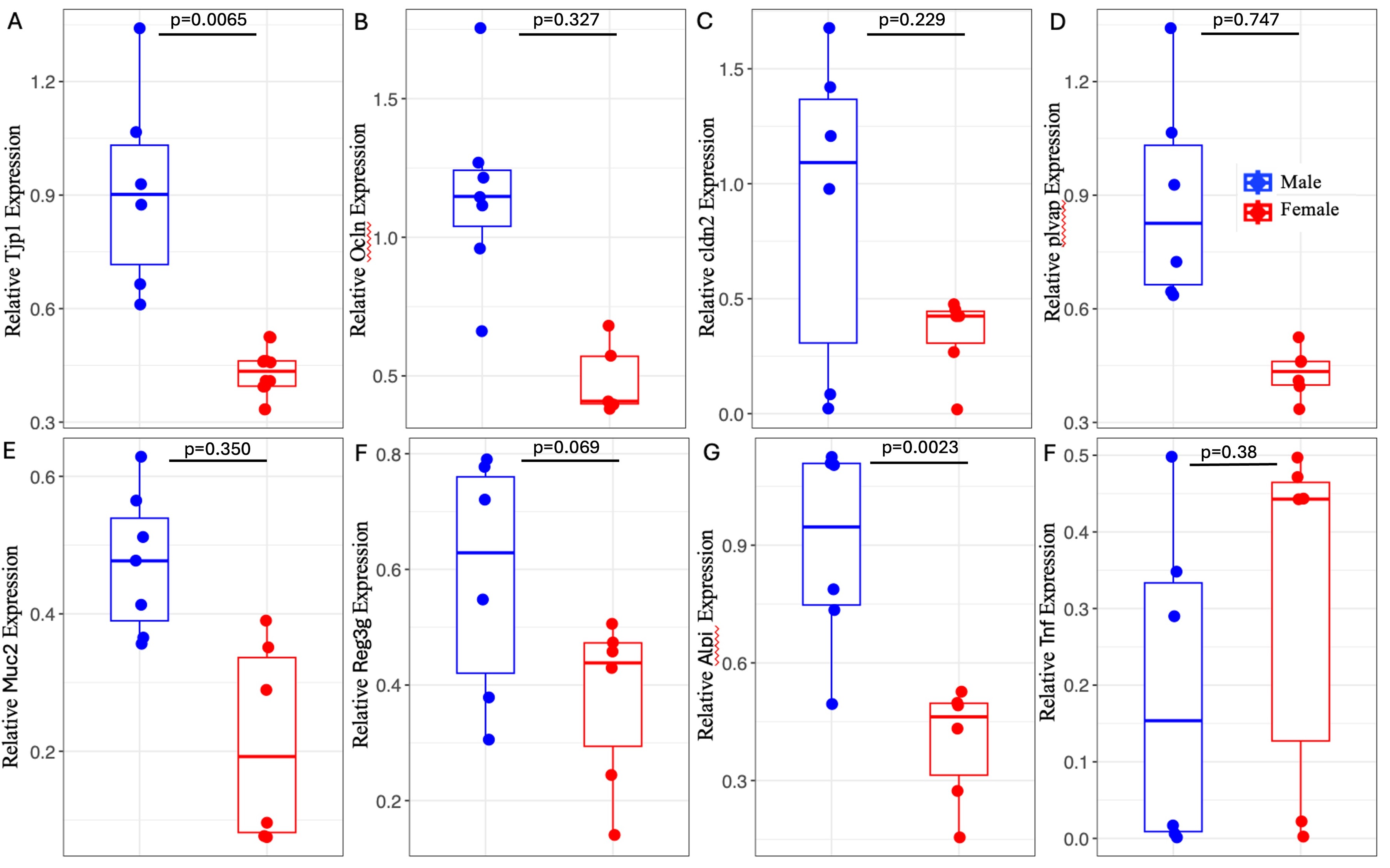

**Supplemental Figure 11 | Transcriptional markers of intestinal barrier structure, mucosal defense, and inflammatory tone across KO males and females**

Relative expression of genes spanning multiple functional layers of the intestinal barrier in KO male and KO female mice. (A) *Tjp1* and (B) *Ocln*, core tight junction–associated proteins; (C) *cldn2*, a pore-forming tight junction component; (D) *plvap*, an endothelial permeability marker; (E) *Muc2*, a mucus layer structural component; (F) *Reg3g*, an epithelial antimicrobial peptide; (G) *Alpi*, an anti-inflammatory epithelial enzyme; and (H) *Tnf*, a pro-inflammatory cytokine. Box-and-whisker plots depict the median (center line), interquartile range (box), and full data range (whiskers), with individual points representing independent biological replicates. Blue points/boxes represent KO males, and red points/boxes represent KO males. Statistical analyses were selected based on Shapiro–Wilk normality and homogeneity of variance testing: Welch’s t test for panel A; unpaired two-tailed Student’s t tests were applied for panels B, E,F,G ; and Mann–Whitney U tests for panels C,D,H. Each group included 5–7 independent biological replicates (n = 5–7 per group). Exact P values are indicated in the respective panels.

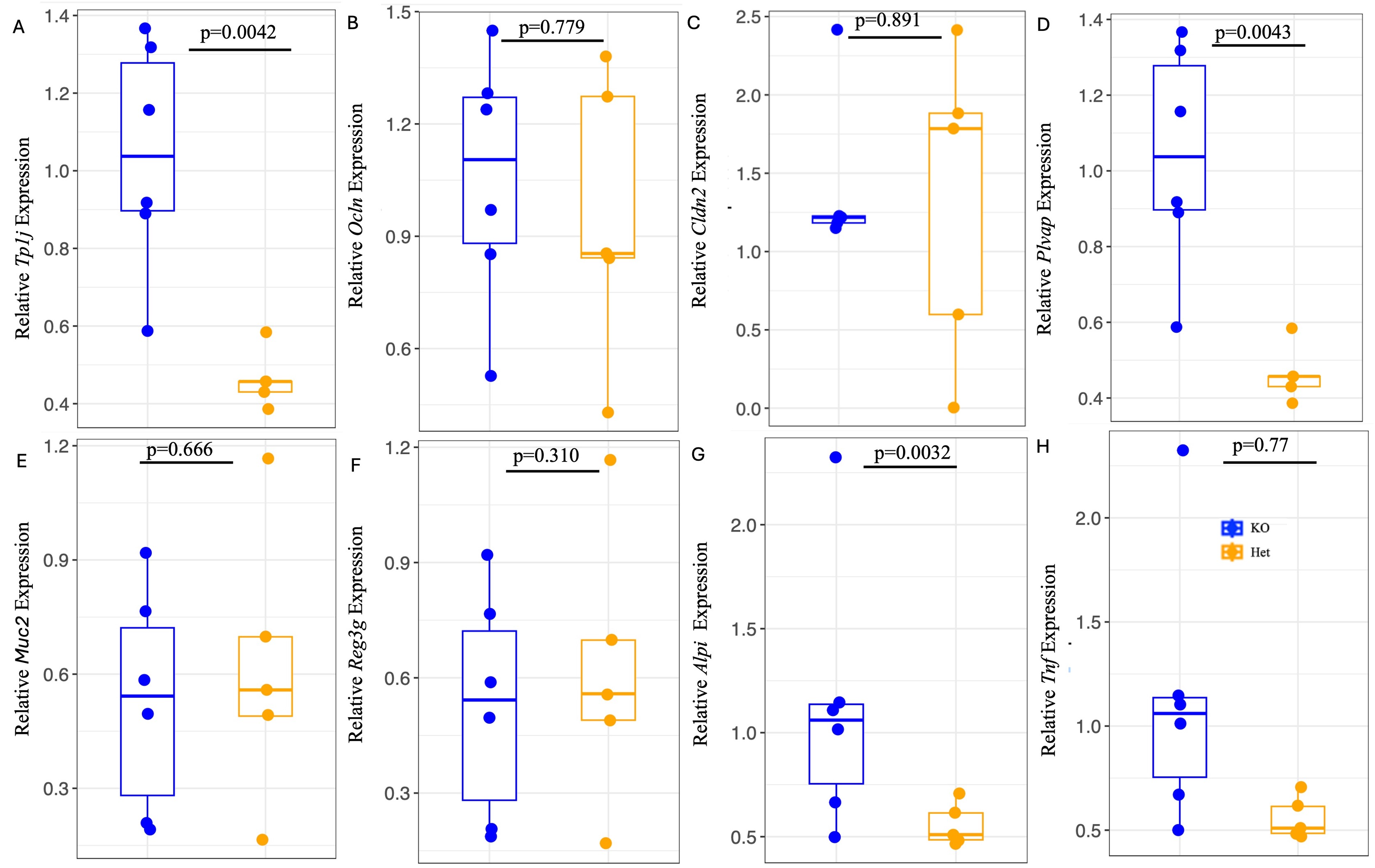

**Supplemental Figure 12 | Transcriptional markers of intestinal barrier structure, mucosal defense, and inflammatory tone across KO males and Heterozygous (Het) females**

Relative expression of genes spanning multiple functional layers of the intestinal barrier in KO male and Het female mice. (A) *Tjp1* and (B) *Ocln*, core tight junction–associated proteins; (C) *cldn2*, a pore-forming tight junction component; (D) *plvap*, an endothelial permeability marker; (E) *Muc2*, a mucus layer structural component; (F) *Reg3g*, an epithelial antimicrobial peptide; (G) *Alpi*, an anti-inflammatory epithelial enzyme; and (H) *Tnf*, a pro-inflammatory cytokine. Box-and-whisker plots depict the median (center line), interquartile range (box), and full data range (whiskers), with individual points representing independent biological replicates. Blue points/boxes represent KO males, and orange points/boxes represent heterozygous (Het) females. Statistical analyses were selected based on Shapiro–Wilk normality and homogeneity of variance testing: Welch’s t test for panel A; unpaired two-tailed Student’s t tests were applied for panels B, E,F,G ; and Mann–Whitney U tests for panels C,D,H. Each group included 5–7 independent biological replicates (n = 5–7 per group). Exact P values are indicated in the respective panels.

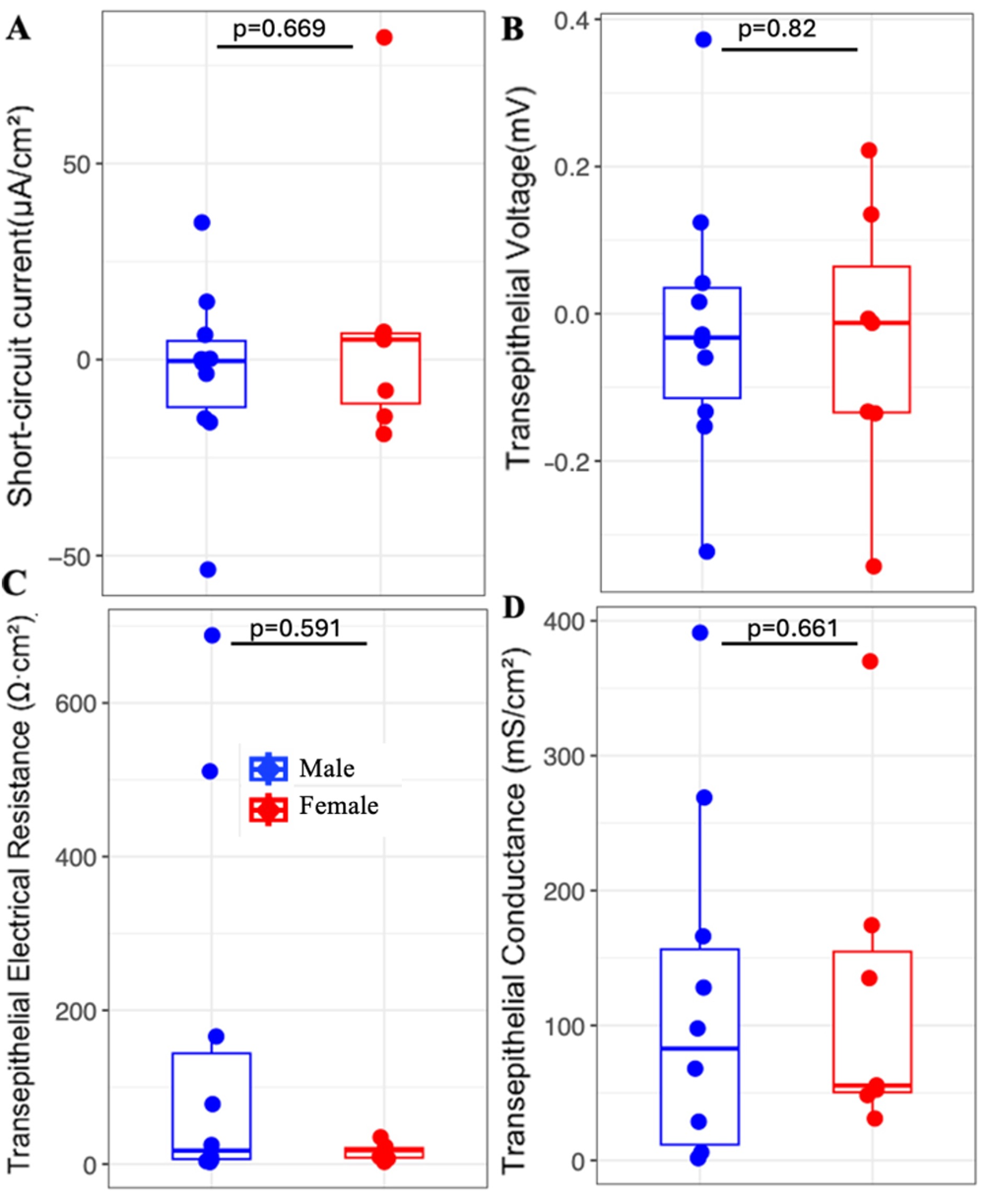
**Extended Data Figure 13 | Sex-associated differences in intestinal epithelial barrier physiology in WT mice.**
Ex vivo assessment of intestinal epithelial barrier function in WT male and WT female mice using Ussing chamber electrophysiology. (A) Short-circuit current (I_sc_), (B) transepithelial voltage (V_t_), (C) transepithelial electrical resistance (TER), and (D) transepithelial conductance (G_t_). Box-and-whisker plots depict the median (center line), interquartile range (box), and full data range (whiskers), with individual points representing independent biological replicates. Blue points/boxes represent WT males, and red points/boxes represent WT females. Statistical analyses were selected based on Shapiro–Wilk normality and homogeneity of variance testing: an unpaired two-tailed Student’s *t* test was applied for panel B, while Mann–Whitney *U* tests were used for panels A, C, and D. Each group included 5–9 independent biological replicates (*n* = 5–9 per group). Exact *P* values are indicated in the respective panels.

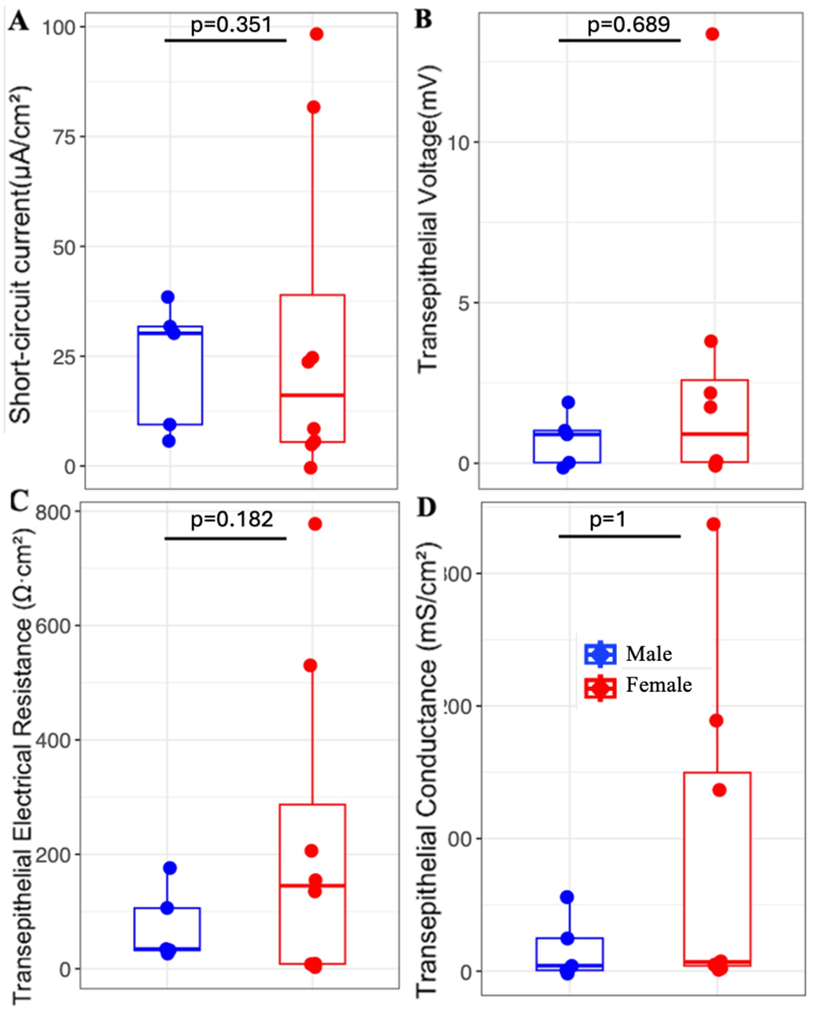
 **Supplemental Figure 14 | Sex-associated differences in intestinal epithelial barrier physiology in KO mice.**
Ex vivo assessment of intestinal epithelial barrier function in KO male and KO female mice using Ussing chamber electrophysiology. (A) Short-circuit current (I_sc_), (B) transepithelial voltage (V_t_), (C) transepithelial electrical resistance (TER), and (D) transepithelial conductance (G_t_). Box-and-whisker plots depict the median (center line), interquartile range (box), and full data range (whiskers), with individual points representing independent biological replicates. Blue points/boxes represent KO males, and red points represent KO females. Statistical analyses were selected based on Shapiro–Wilk normality and homogeneity of variance testing: Mann–Whitney *U* tests were used for panels A, B, C, and D. Each group included 5–9 independent biological replicates (*n* = 5–9 per group). Exact *P* values are indicated in the respective panels.

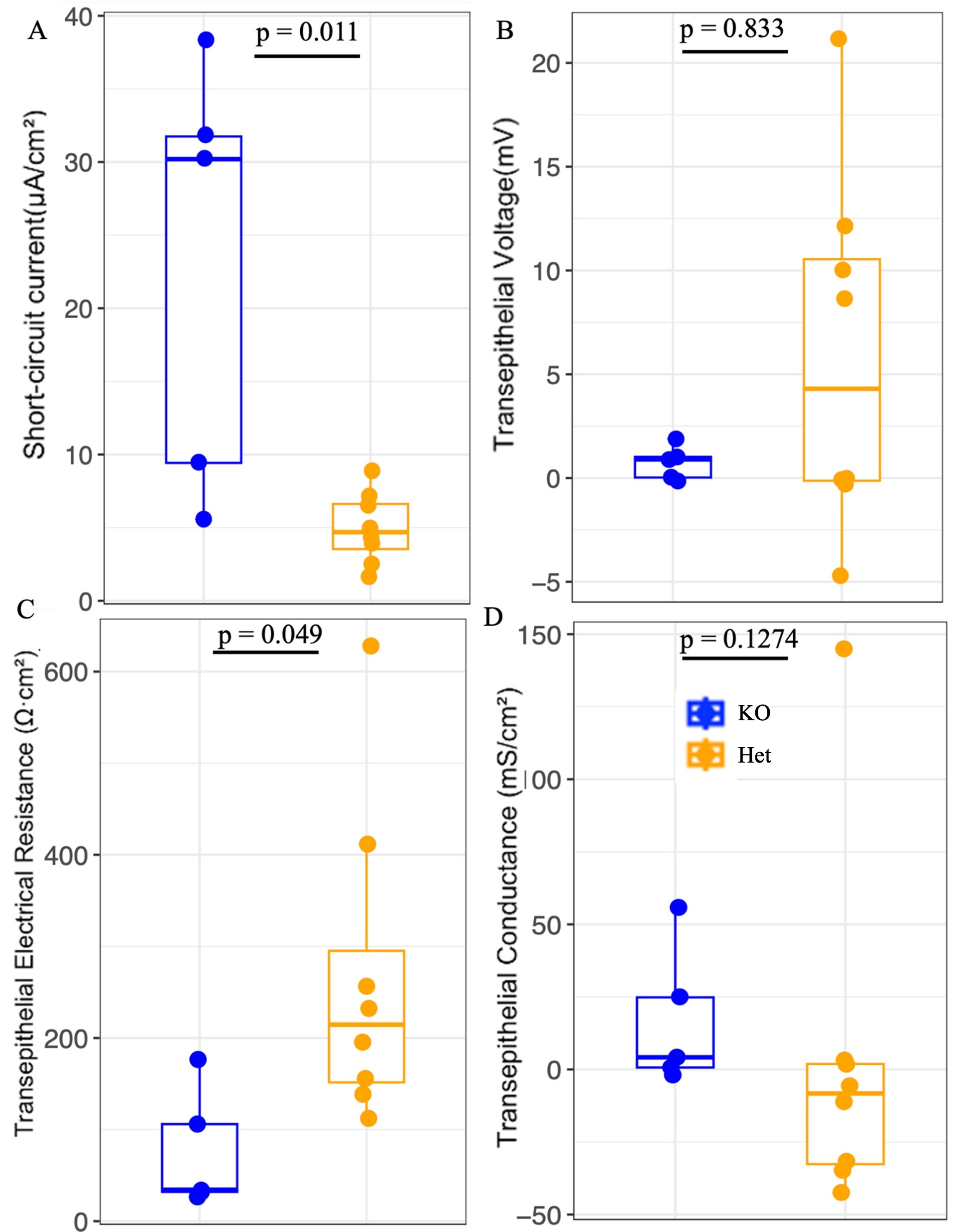

**Extended Data Figure 15 | Differences in intestinal epithelial barrier physiology in KO male and Heterozygous (Het) female mice.**
Ex vivo assessment of intestinal epithelial barrier function in KO male and Het female mice using Ussing chamber electrophysiology. (A) Short-circuit current (I_sc_), (B) transepithelial voltage (V_t_), (C) transepithelial electrical resistance (TER), and (D) transepithelial conductance (G_t_). Box-and-whisker plots depict the median (center line), interquartile range (box), and full data range (whiskers), with individual points representing independent biological replicates. Blue points/boxes represent KO males, and orange points represent KO females. Statistical analyses were selected based on Shapiro–Wilk normality and homogeneity of variance testing: Mann–Whitney *U* tests were used for panels A, B, C, and D. Each group included 5–9 independent biological replicates (*n* = 5–9 per group). Exact
